# Response to azacitidine and the TIM-3 inhibitor, sabatolimab, in MDS and CMML is associated with distinct immune and antigen-presentation states

**DOI:** 10.64898/2026.11.04.764360

**Authors:** Aonkon Dey, Neal P. Smith, Alice Tirard, Benjamin Arnold, Thomas Chan, Jennifer Lombardi-Story, Timothy Casson, Tanya T. Behnan, Sergio Aguilar-Fernandez, Amir T. Fathi, Gabriela S. Hobbs, Hanno Hock, Philip C. Amrein, Patrick Reeves, Max Jan, Lloyd Bod, Timothy A. Graubert, Andrew M. Brunner, Rupa Narayan, Alexandra-Chloé Villani

**Affiliations:** Center for Immunology and Inflammatory Diseases, Massachusetts General Hospital, Boston, MA; Krantz Family Center for Cancer Research, Massachusetts General Hospital, Boston, MA; Broad Institute of Massachusetts Institute of Technology and Harvard, Cambridge, MA; Mass General Brigham Cancer Institute, Boston, MA; Harvard Medical School, Boston, MA; Division of Hematology/Oncology, Massachusetts General Hospital, Boston, MA; Vaccine and Immunotherapy Center, Massachusetts General Hospital, Boston, MA

**Keywords:** MDS, CMML, scRNAseq, HSPC, HSC, LSPC, TIM3 or TIM-3, hypomethylating agents, HMA, azacitidine, sabatolimab

## Abstract

Targeting the immune checkpoint TIM-3 is of interest in myeloid neoplasms, yet the hematopoietic and immune changes associated with anti–TIM-3 therapy remain poorly characterized. We used single-cell RNA sequencing to profile bone marrow (BM) and peripheral blood from patients with myelodysplastic syndrome (MDS) and chronic myelomonocytic leukemia (CMML) treated with the hypomethylating agent azacitidine combined with sabatolimab, a monoclonal antibody targeting TIM-3, and integrated Human Cell Atlas reference datasets for comparison. CITE-seq profiling showed that TIM-3 was predominantly expressed on dendritic cells and NK cells, with variable expression across other populations, including CD34 cells and CD8 T cells. At baseline, we identified a population of CD34 myeloid-biased progenitors specific to MDS/CMML samples (“HSPC-like”) characterized by inflammatory gene expression programs and features of impaired differentiation. In MDS, elevated baseline HLA class II expression across hematopoietic stem and progenitor cells (HSPCs), HSPC-like cells, and multiple myeloid populations was associated with response. Consistent with this finding, trajectory analysis suggested that nonresponders bypassed HLA class II-high states during myeloid differentiation. In MDS, the presence of distinct CD8 T-cell populations and increased interferon-response programs in CD8 T and NK cells at baseline was also associated with response. In both MDS and CMML, treatment among responders was associated with contraction of BM CD8 and CD4 T-cell populations characterized by type I interferon-responsive and inflammatory programs, including *IFIT1, IFIT2,* and *TNF*, along with expansion of a distinct *GZMH/IFITM1*-expressing cytotoxic CD8 T-cell population. Together, these findings identify baseline antigen-presentation and immune-response programs associated with therapeutic response and reveal coordinated remodeling of myeloid and lymphoid compartments during azacitidine/sabatolimab treatment in MDS and CMML.

## INTRODUCTION

Myelodysplastic syndrome (MDS) and chronic myelomonocytic leukemia (CMML) are clonal hematologic malignancies characterized by progressive bone marrow failure and risk of progression to acute myeloid leukemia (AML). While hypomethylating agents (HMAs), such as decitabine or azacitidine, are commonly used as first line therapy^1–5^, median overall survival (mOS) of patients with MDS and CMML remains poor, particularly for patients who do not initially respond to HMA therapy or who subsequently lose treatment response^6–9^.

Several lines of investigation have provided a rationale for combining HMAs with immunotherapy. HMA therapy *in vitro* and *in vivo* has been shown to induce inflammatory and immune-related gene expression programs^10,11^, including interferon response^12^, reactivation of endogenous retroviral elements^13,14^, and antiviral responses^15^. HMA therapy may also upregulate immune checkpoints, including PD-L1, PD-L2, and CTLA4^16^, which may be a potential resistance mechanism^17,18^.

The immune checkpoint protein TIM-3 has emerged as a target of interest in myeloid neoplasms, in part because TIM-3 is expressed on leukemic stem cells (LSCs)^19–22^, and treatment with anti-TIM-3 antibodies in xenograft models has been shown to reduce LSC engraftment^20^. Elevated TIM-3 expression has also been associated with MDS disease progression^23^ ^21^ and observed in bone marrow (BM) CD4 T cells of patients with AML relapsing following allogeneic hematopoietic stem cell transplantation (allo-HSCT) ^24^. Phosphotidylserine (PtdSer), GAL-9, and HMGB-1 have been described among putative modulators of TIM-3 function^25^. TIM-3 expression has also been observed on CD4 Th1 cells, CD8 cytotoxic T cells and CD11b cells,^26,27^ with studies suggesting that TIM-3 negatively regulates Th1 cells and their expansion^27,28^. Significant TIM-3 expression has also been reported in natural killer (NK), myelomonocytic, and dendritic cells (DCs)^26,29–31^, with TIM-3 down-regulation observed during DC maturation and activation^29^. Furthermore, conditional knockout of TIM-3 in DCs was shown to promote anti-tumor immunity^32^.

Sabatolimab (MBG453) is an IgG4 monoclonal antibody targeting TIM-3^33,34^*. In vitro* studies have shown that sabatolimab can mediate antibody-dependent cellular phagocytosis (ADCP), enhance T cell killing of TIM-3 expressing neoplastic cells, promote cytokine secretion by DCs, and inhibit GAL-9 or PtdSer binding to TIM-3^34^. Sabatolimab has been evaluated as monotherapy and in combination with decitabine or azacitidine, in patients with high risk (HR)/very high risk (vHR)-MDS, CMML and AML in a phase 1b study (NCT03066648)^35,36^. Among evaluable patients, the overall response rate (ORR) to sabatolimab combined with HMAs was 56.9% and 66.7% in patients with HR/vHR-MDS and CMML, respectively, with a mOS of 26.7 months^35^. In randomized phase III data comparing azacitidine plus sabatolimab with azacitidine plus placebo in patients with higher-risk MDS or CMML-2, the mOS was similar between the groups (22.3 vs 18.8 months, p=0.0825), and the study did not meet the significance threshold (NCT04266301)^37^. The safety profile of HMA/sabatolimab was generally been found to be comparable to that of HMA/placebo, with adverse events primarily related to myelosuppression and with rare severe immune-related adverse events^38^. While there was a lack of a significant survival benefit in this specific phase III study, TIM-3 and its putative ligands (including GAL-9) remain potential targets of interest in myeloid neoplasms, given the distinct biology and expression profile of TIM-3, including in LSCs, DCs and myelomonocytic populations.

Given the limited understanding of the immunological effects associated with targeting TIM-3 in combination with HMA therapy in patients with myeloid malignancies, we performed single cell RNA sequencing (scRNAseq) of BM and peripheral blood mononuclear cells (PBMCs) from subjects with MDS and CMML treated at our institution with sabatolimab-based therapy on NCT03066648. We integrated these data with publicly available scRNAseq data of BM and PBMCs of healthy donors from the human cell atlas (HCA)^39^, jointly analyzing over 800,000 single cell profiles. We sought to characterize disease-associated hematopoietic and immune states, define cellular and transcriptional changes associated with azacitidine/sabatolimab combination therapy, and identify baseline cellular and transcriptional features associated with therapeutic response.

## RESULTS

### Cohort overview and single-cell study design

We generated scRNA-seq on BM and PBMC samples from 16 patients diagnosed with either de novo MDS (n=7), CMML (n=5), or relapsed/refractory MDS (RR-MDS; n=4) **(Fig. 1A, Supplemental tables 1-3)**. Twelve patients had received the anti-TIM-3 antibody (sabatolimab), administered either as monotherapy (n=3, RR-MDS), in combination with the anti-PD-1 antibody spartalizumab (n=1, RR-MDS), or with azacitidine (n=5, de-novo MDS; n=3, CMML) as part of a phase 1b clinical trial. The remaining 4 patients had been treated with HMA therapy alone as part of standard care (n=2, de novo MDS; n=2, CMML).

**Figure 1.**
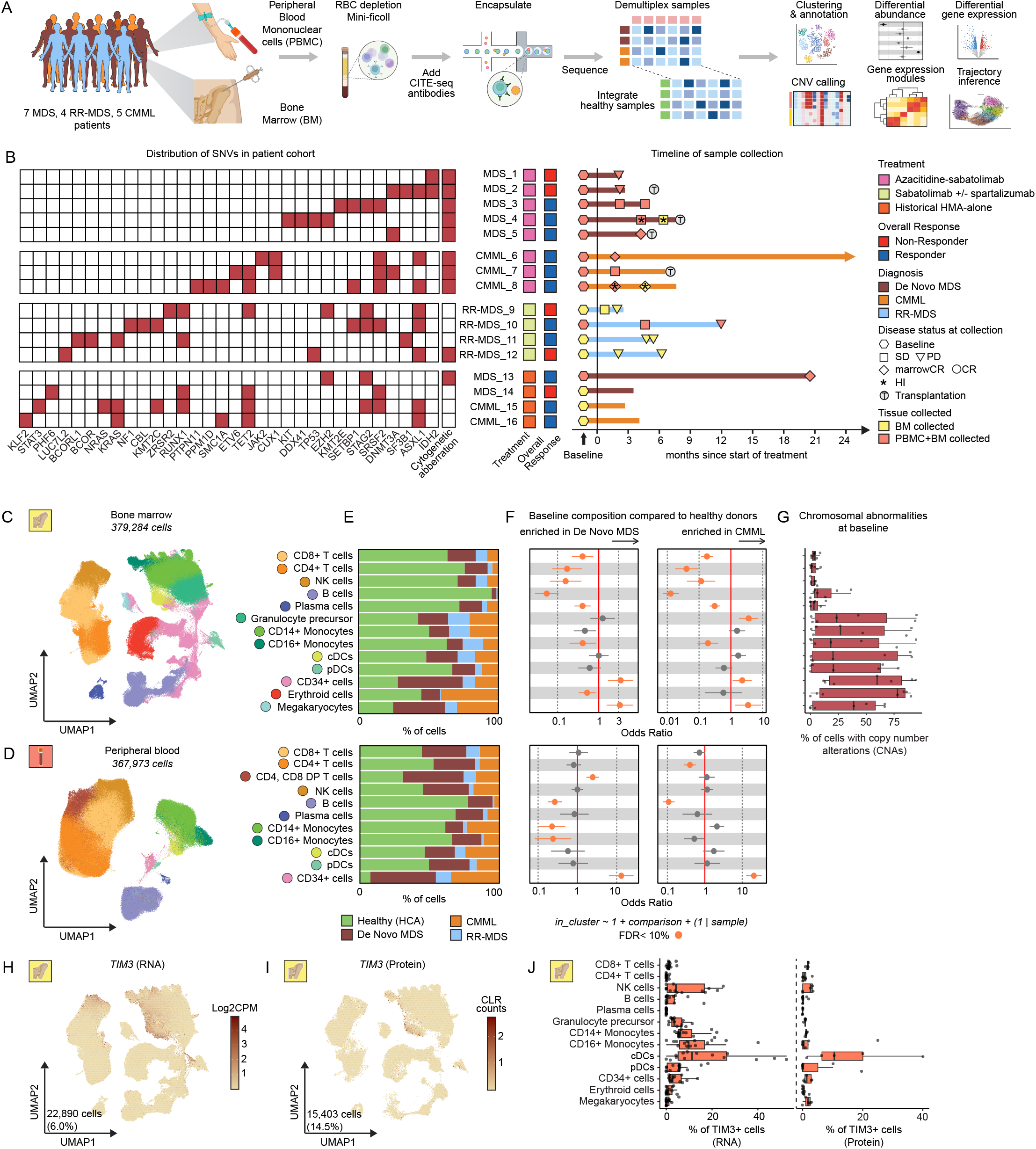
A. Study design and single-cell multiomic landscape of bone marrow and peripheral blood in de novo MDS and CMML at baseline. Experimental design schematic. **B.** Heatmap showing the variant allele frequency (VAF) of the most abundant single nucleotide variants (SNVs) in recurrently mutated gene loci across individual patients at baseline. **C-D.** Uniform manifold approximation and projection (UMAP) embedding of (**C**) 379,284 cells from bone marrow (BM) and (**D**) 367,973 cells from peripheral blood (PB) colored by major cell lineage. **E.** Cell composition for each lineage by origin from patient groups and healthy donor cells from the human cell atlas (HCA). Bars represent the fraction of cells in each lineage derived from the corresponding group in BM (top) and PBMC (bottom). **F.** Cell lineage abundance analysis comparing baseline samples from de novo MDS (left panels) and CMML (right panels) to healthy donor cells in the BM (top panels) and PBMC (bottom panels). Points represent odds-ratio with error bars representing 95% confidence intervals. Color indicates FDR < 0.1 (logistic regression). **G.** Chromosomal abnormalities at baseline by major cell lineage. Dots represent the percent of cells from each donor in each lineage with NUMBAT inferred copy number variation (CNV) at predefined loci. **H-I**. UMAP feature plots using color to indicate RNA transcript (left; log2CPM) and protein (right; CLR) levels of *HAVCR2*/TIM3. Number and percentage correspond to cells with non-zero expression. **J.** Percent of cells from each donor by cell lineage with non-zero counts for TIM3 RNA and TIM3 protein after background signature removal. In **G** and **J** box plots show median (line) and IQR, with whiskers no more than 1.5x IQR.

After quality control filtering, pretreatment BM baseline samples from 14 patients were available for downstream analysis. Post-treatment samples were variably available across timepoints, including early (n=5), late (n=4), or end of treatment (EOT; n=9) collections. Paired PBMC samples were included where available (**Supplemental table 1**). CITE-seq data were generated for a subset of samples where feasible.

The cohort captures patients harboring common recurrent mutations observed in MDS and CMML **(Fig. 1B)**. For this analysis, patients who achieved stable disease (SD) or better as best response were classified as overall responders (**Fig. 1B**). Among the 8 patients treated with sabatolimab combined with azacitidine, 2 were non-responders and 6 were responders, including all three CMML patients. Disease status at each sampling timepoint is indicated in **Fig. 1B**.

To enable robust clustering and comparative gene expression analyses, single cell RNAseq datasets from our study cohort were integrated with healthy donor BM and PBMC datasets from the HCA repository^39^. In total, 206,183 BM and 193,347 PBMC cells from our cohort were integrated with 240,650 BM and 200,000 PBMC cells from 8 healthy donors. Following doublet removal and low-quality subcluster filtering, 379,284 BM and 367,973 PBMC cells were retained for downstream analysis (**Fig. 1C-D, Supplementary Fig. 1A-B**).

Within the BM compartment, we identified 13 major hematopoietic lineages: CD8 T cells (*CD3D, CD8A*), CD4 T cells (*CD3D, CD4*), natural killer (NK) cells (*NCAM1*), B cells (*MS4A1*), plasma cells (*JCHAIN*), granulocyte precursors (*MPO, AZU1*), CD14 monocytes (*CD14*), CD16 monocytes (*FCGR3A*), conventional dendritic cells (cDCs) (*CD1C, CLEC10A*), plasmacytoid dendritic cells (pDCs) (*JCHAIN, IL3RA*), CD34 expressing cells (*CD34*), erythroid cells (*HBB, AHSP*) and megakaryocyte lineage cells (*PF4, PPBP*) (**Fig. 1E, top panel; Supplementary Fig. 1C, Supplemental table 4-5**). In PBMC, 11 major lineages were detected: CD8 T cells, CD4 T cells, CD4/CD8 double positive T cells, NK cells, B cells, plasma cells, CD14 monocytes, CD16 monocytes, cDCs, pDCs, and CD34 cells (**Fig. 1E, bottom panel; Supplementary Fig. 1D, Supplemental table 4-5**).

We first compared the relative lineage abundance between healthy donors and patients with de novo MDS or CMML using pretreatment samples (**Fig.1F, Supplemental table 6**). Patients exhibited reduced frequencies of BM CD8 and CD4 T and NK cells relative to healthy donors (**Fig. 1F**), while B cell frequencies were markedly decreased in both the BM and PBMC compartments. Monocytic populations showed disease-specific alterations; CD14 and CD16 monocytes were significantly reduced in PBMCs in de novo MDS (CD14: OR=0.007, p=0.018; CD16: OR=0.016, p=0.03), and CD16 monocytes were reduced in BM of both de novo MDS (OR=0.19, p=0.0018) as well as CMML (OR=0.072, p= 0.0013). In contrast, there was enrichment of granulocyte precursors in the BM of CMML samples (OR= 3.62, p= 0.0044). Both MDS and CMML were associated with increased frequencies of CD34 cells in BM (MDS: OR=3.39, p=0.017; CMML: OR= 2.28, p=0.072) and PBMCs (MDS: OR=13, p=0.0001; CMML: OR=20.4, p=0.0000016) compared to healthy donors. Copy number alterations, inferred from single-cell CNV analysis, were detected across the CD34, myelomonocytic, megakaryocyte, and erythroid lineages (**Fig. 1G, Supplementary Fig. 1E**), consistent with clonal hematopoiesis described in MDS and _CMML_40–42.

Given that sabatolimab targets TIM-3, we next assessed TIM-3 (*HAVCR2*) expression at both transcript (n=14) and surface protein (n=5) expression levels across cell lineages to identify potential cellular modulators of therapeutic response (**Fig.1H-J, Supplementary Fig.1F-H**). Conventional DCs had the highest median TIM-3 protein expression, with additional expression observed in NK cells, monocytes, and CD34 cells. The TIM-3 ligand, HMGB1 (*HMGB1), was* highly expressed in CD34 and erythroid cells, whereas GAL-9 (*LGALS9*) expression was enriched in granulocyte precursors, monocytes, and DCs (**Supplementary Fig. 1I**).

In summary, this dataset captures the major hematopoietic lineages across BM and PBMCs, with data revealing baseline abundance differences in cell lineages involved in acquired immunity (T and B cells) as well as innate immunity (NK cells, CD16 monocytic subset) along with left shifted hematopoiesis with CD34 enrichment in MDS/CMML. Expression of TIM-3 and its ligands were identified across CD34 cells and cells involved in antigen presentation (DCs, monocytes) and/or innate immune functions (NK cells, monocytes).

### CD34 hematopoietic stem and progenitor-like cells in MDS and CMML shift toward a dysplastic and inflammatory state

To further characterize disease-associated alterations in MDS and CMML, we performed sub-clustering of the CD34 **(Fig. 2A-E, Supplementary Fig. 2A-B)**, myelomonocytic-DC **(Fig. 2F-J)**, and erythroid-megakaryocyte compartments respectively (**Supplementary Fig. 3A-B)**.

**Figure 2.**
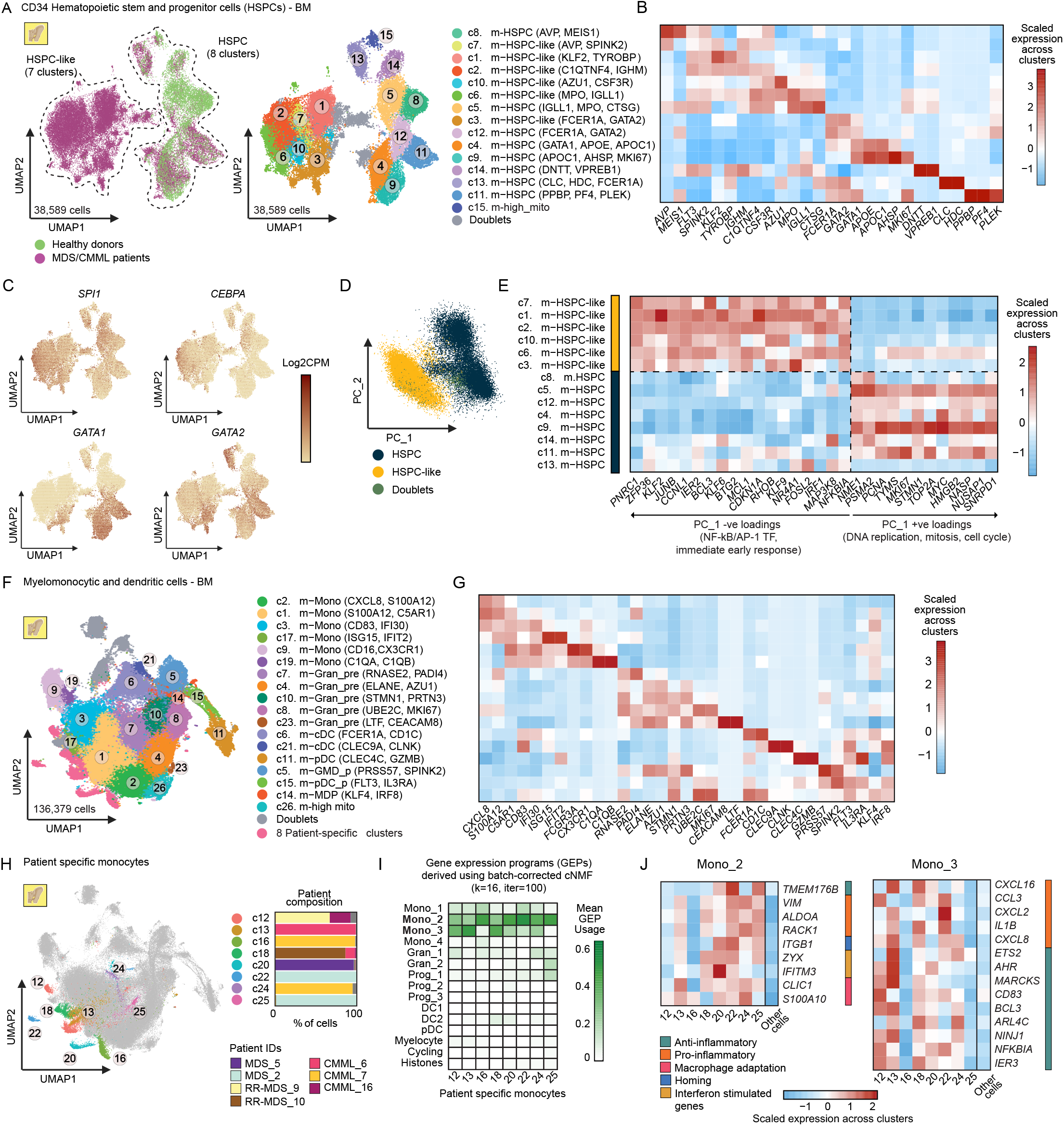
Subclustering of HSPCs, and myelomonocytic and dendritic cells in bone marrow reveals dysplastic and inflammatory HSPC state and patient-specific monocyte populations. **A.** UMAP embedding of 38,589 HSPCs from BM colored by overall cell origin group (left, healthy donors or MDS/CMML patients) and subcluster (right) annotations. **B.** Heatmap showing discriminative markers for each cluster in **A** when compared with every other cluster. Color scale denotes the normalized gene expression (mean zero, unit variance) across cell clusters. **C.** UMAP feature plots of myeloid-primed MPP markers using color to indicate RNA transcript levels (log2CPM). **D.** Dotplot of first and second principal component weights of all HSPC cells colored by subcluster groups, HSPC and HSPC-like. **E.** Heatmap showing markers with positive and negative loadings in the first principal component. Color scale denotes the normalized gene expression (mean zero, unit variance) across cell clusters. **F.** UMAP embedding of 136,379 myelomonocytic and dendritic cells (DCs) from BM colored by subcluster annotations. **G.** Heatmap showing discriminative markers for each cluster in **F** when compared to every other cluster. Color scale denotes the normalized gene expression (mean zero, unit variance) across cell clusters. **H.** UMAP embedding showing patient-specific myelomonocytic and DC clusters (left) and patient-specific cluster composition (right). Bars represent the fraction of cells in each cluster that were derived from the corresponding patient. **I.** Heatmap showing mean usage of gene expression programs derived from cNMF in patient-specific myelomonocytic and DC clusters. **J.** Heatmap showing select genes from the Mono_2 (left) and Mono_3 (right) gene expression programs across patient-specific clusters. Color scale denotes the normalized gene expression (mean zero, unit variance) across cell clusters.

Subclustering of 38,589 BM CD34 cells identified two major subsets: a shared subset comprising cells from both healthy donors and patients, defined as hematopoietic stem and progenitor cells (HSPCs), and a second subset composed exclusively of patient-derived cells, termed HSPC-like based on their transcriptional profiles (**Fig. 2A**). The HSPC subset encompassed canonical stem cell and progenitor populations, including hematopoietic stem cells (c8; *AVP*) and multipotent progenitors (MPPs) (c5, c14; *FLT3* and *SPINK2*) (**Fig. 2B, Supplemental tables 4-5)**. Within the MPP compartment, we observed lineage priming toward myeloid fate (c5; *MPO*, *AZU1*, and *CSF3R*) and lymphoid fate (c14; *DNTT* and *VPREB1*). Additional progenitor populations included megakaryocyte-erythroid progenitors (MEPs; c12; *GATA1*, *GATA2*, and *FCER1A*), megakaryocyte-biased progenitors (c11; *PPBP*, *PF4*), erythroid-biased progenitors (c4, c9; *APOE* and *APOC1*), and mast cell/eosinophil/basophil-biased progenitors (c13; *CLC*, *HDC*).

While the HSPC-like subset expressed gene programs resembling stem and progenitor states-including stem-like (c7), MPP-like (c1, c2, c6, c10), and MEP-like (c3) signatures-there was a pronounced skew towards myeloid priming, as indicated by the increased expression of *SPI1, CEBPA, GATA1* and *GATA2* **(Fig. 2C)**.

To further define the transcriptional divergence between the HSPC and HSPC-like subsets, we examined the principal components underlying the embedding of BM CD34 cells. The first principal component (PC1) captured the primary axis of separation between HSPCs and HSPC-like cells (**Fig. 2D**). Gene loading analysis revealed two opposing transcriptional programs independent of lineage identity. HSPCs were enriched for genes associated with DNA replication (*NME1. PCNA, TYMS, TOP2A*), mitosis (*STMN1, HMGB2, PSMA2*) and cell cycle regulation (*MKI67, NASP, SNRPD1*), consistent with a cycling state (**Fig. 2E**). In contrast, HSPC-like cells upregulated inflammatory and stress-response programs, including transcription factors and signaling regulators (*NFKBIA, FOSL2, IRF1, MAP3K8*, and *BCL3*), immediate early genes (*IER2, JUNB, NR4A1, BTG2*, *and CDKN1A*), post-transcriptional regulators (*ZFP36* and *CCNL1*), and stress-associated KLF family transcription factors (*KLF2*, *KLF6*, *KLF9*).

Collectively, these findings identify a distinct population of CD34 HSPC-like cells in MDS and CMML characterized by inflammatory signaling and stress-response programs (**Fig. 2E**). Importantly, no significant differences in quality control metrics were observed between HSPC and HSPC-like cells (**Supplementary Fig. 2A-B**).

### Shared gene programs between patient-specific monocytes consist of early activation, inflammation, and adaptation

After re-clustering 136,379 myelomonocytic and dendritic cells, we observed five major sublineages shared between healthy donors and patients with MDS or CMML, namely classical monocytes (c2, c1, c3, c17; *CD14* and *FCN1*), non-classical monocytes (c9, c19; *CD16*), granulocyte precursors (c8, c10, c4, c7; *ELANE, AZU1, MPO*), cDCs (c6, c21; *CD1C*) and pDCs (c11; *IL3RA*) (**Fig. 2F-G, Supplemental tables 4-5**). There were also a number of clusters suggestive of progenitors with lineage bias (c5, c14, c15) co-expressing various levels of *CD34, PRSS57, SPINK2, FLT3* and cycling markers.

We also identified a number of patient specific monocyte clusters (**Fig. 2H, Supplementary Fig. 2C**) that were transcriptionally distinct but lacked clear marker genes (**Supplementary Fig. 2D-E**). These clusters were robust to integration techniques (**Supplementary Fig. 2F-H**). Hypothesizing the presence of shared gene programs unrelated to cell type identity driving the specificity of these clusters, we used consensus non-negative matrix factorisation to derive 16 stable programs (**Supplementary Fig. 2I-K, Supplemental table 7**). Two such programs, Mono_2 and Mono_3, had the highest usage by patient-specific monocyte clusters (**Fig. 2I**). Examining the genes comprising such programs, we observe genes related to early activation, inflammation, and adaptation (**Fig. 2J**). Combinatorial usage of these programs and the monocyte identity programs captures the transcriptional heterogeneity driving these clusters (**Supplementary Fig. 2L**). In baseline samples, we observed Mono_2 usage enriched in patients who eventually had progressive disease (PD) and Mono_3 usage enriched in patients with CMML who eventually achieved a response, with healthy donors having the lowest usage of either program (**Supplementary Fig. 2M**).

After re-clustering 26,523 erythroid cells and megakaryocytes, we identified two subsets of megakaryocytes (c6, c7; *ITGA2B, PLEK, PF4*) and five subsets of erythroid cells, defined by *HBB, HBA1, AHSP* **(Supplementary Fig. 3A-B, Supplemental tables 4-5**). Among megakaryocyte subsets, c7 expressed TUBB1, a late-stage marker of mature megakaryocytes. Erythroid subsets were marked by increasing expression of *GYPA* and decreasing expression of *KIT* and *APOE*, suggestive of maturation stages in the order of c4, c1, c2 and c5. Cluster 3 (c3; *ALYREF* and *NME2*) appears to be an early stage erythroid cell subset specific to myeloid malignancy (**Supplementary Fig. 3B**).

Thus, in addition to shared major myelomonocytic lineages between healthy donors and patients, we observed patient specific monocytes clusters characterized by shared gene expression programs involved in activation and inflammation, with data suggesting associations between program usage and eventual response.

### Effects of combinatorial sabatolimab/azacitidine therapy on progenitor and myelo- monocytic compartments in MDS and CMML

Upon re-clustering to define granular cell subsets, we next evaluated the effects of combined sabatolimab and azacitidine therapy. Analyses focused on the BM compartment due to the availability of paired pre- and post-treatment samples from responders with de novo MDS or CMML treated with combination therapy. Owing to limited sample size among non-responders (n=2), comparative analyses of treatment-associated changes in this group were not feasible. The mean post-treatment sampling interval was 115 days for de novo MDS and 56 days for CMML (**Fig. 3A**).

**Figure 3.**
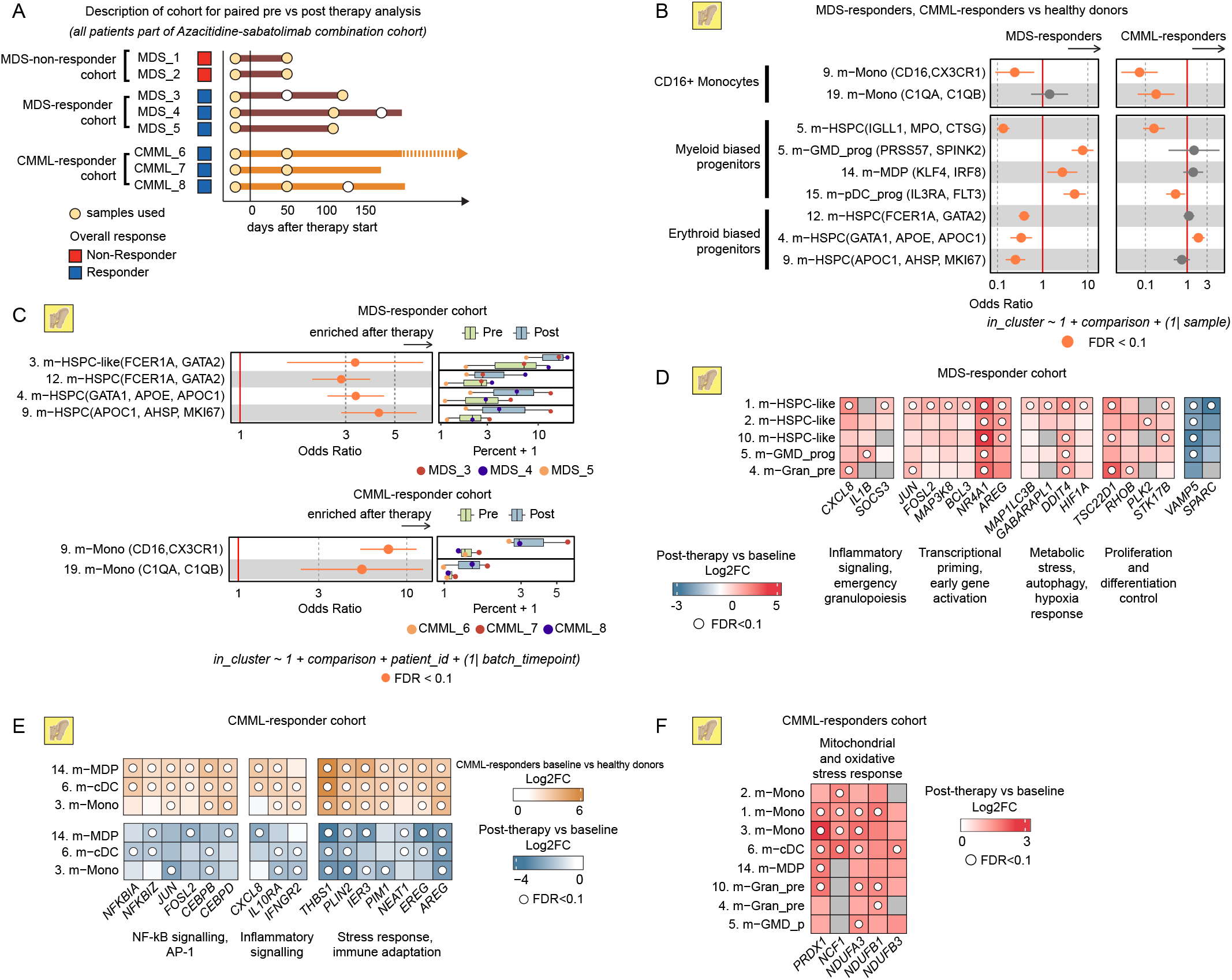
Treatment-associated shifts in cell subset abundance and gene expression in myeloid and progenitor compartments of MDS and CMML responders. **A.** Swimmer’s plot showing the pre- and post-treatment sample timepoint distribution from the CMML and de novo MDS cohorts. Yellow dots indicate samples used for comparing pre- and post-treatment samples. **B.** Cell cluster abundance analysis comparing MDS-responders (left) and CMML-responders (right) to healthy donors at baseline in the BM. **C.** Cell cluster abundance analysis comparing the pre- and post-treatment timepoints for MDS-responders (top) and CMML-responders (bottom) in the BM. **D.** Heatmap showing select differentially expressed genes (DEGs) in MDS-responders comparing post-treatment to baseline samples in the HSPC-like, GMD progenitor and granulocyte precursor clusters in BM. **E.** Heatmap showing select DEGs in specific DC and monocyte clusters in BM among CMML-responders comparing baseline patient samples to healthy donor (top) or post-treatment samples (bottom). **F.** Select DEGs in monocyte, DC and granulocyte precursor clusters in the BM among CMML-responders comparing post-treatment to baseline samples. In **B** and **C**, Points represent odds-ratio with error bars representing 95% confidence intervals. Color indicates FDR < 0.1 (logistic regression). In **C,** box plots show median (line) and IQR, with whiskers no more than 1.5x IQR. In **D-F**, Color scale indicates log2FC and white dots indicate FDR < 0.1.

We first assessed baseline differences in subcluster abundance between healthy donors and pre-treatment patient samples (**Fig. 3B, Supplementary Fig. 4A-B, Supplemental table 6**). In the progenitor compartment, myeloid-primed HPSC-like cells of a dysplastic nature (c1, c2, c6, c7, c10) were highly enriched, consistent with earlier analyses (**Fig. 2A-D**), while MPP (c5) abundance was decreased in both disease groups. In MDS, MEPs and erythroid progenitors were reduced in the progenitor compartment (**Fig. 3B**), whereas granulocyte, monocyte and dendritic cell progenitors (c5, c14, c15) within the myelomonocytic compartment were enriched. Both MDS and CMML samples demonstrated reduced frequencies of CD16 monocytes (c9) (**Fig. 3B**), while inflammatory C1Q,CD16 monocytes (c19) were also reduced in CMML.

Comparison of pre- versus post-treatment samples revealed therapy-associated cellular composition shifts among responders. In CMML, treatment was associated with increased abundance of CD16 monocytes, whereas in MDS, there was an expansion of erythroid progenitors and fully differentiated erythroid populations (**Fig. 3C, Supplementary Fig. 4A**).

Differential gene expression analysis revealed disease-specific transcriptional response to therapy. In MDS, treatment was associated with induction of NF-κB/AP-1 transcription factor signaling, stress adaptation pathways, and apoptotic programs within HPSC-like subsets, granulocyte-monocyte-DC progenitors, and granulocyte precursors (**Fig. 3D, Supplemental table 8**). In CMML, comparison of baseline gene expression to healthy donors demonstrated upregulation of chronic inflammatory stress adaptation programs (*PIM1, IER3, NEAT1*), immunoregulatory signals (*THBS1*), trophic signals (*AREG, EREG*), NF-κB/AP-1, and inflammatory signaling in DCs, DC progenitors and antigen-presenting monocytes (**Fig. 3E, Supplementary Fig. 4C**). These programs were subsequently downregulated in these subsets following therapy. Amongst responders with CMML, we also observed an increase in mitochondrial- and oxidative stress-responsive gene expression across monocytes, DCs and granulocyte precursors post-therapy, suggestive of metabolic rewiring (**Fig. 3F, Supplementary Fig. 4C**).

In summary, amongst responding patients with MDS/CMML, combination sabatolimab/azacitidine therapy was associated with abundance shifts suggestive of disease specific restoration of monocyte and/or erythroid differentiation programs at least in part, while gene expression analysis suggested upregulated inflammatory transcriptional shift in specific progenitor and myeloid precursor subsets in MDS versus downregulation of several chronic inflammatory programs in specific DC/monocytes subsets in CMML. The difference in condition and median post-treatment collection (2 cycles for CMML and 4 cycles in MDS) may have contributed to the distinct transcriptional responses observed between these disease groups.

### Effects of combinatorial azacitidine/sabatolimab therapy on adaptive immunity and T cell states in MDS and CMML

To characterize adaptive immune alterations associated with combined azacitidine and sabatolimab therapy, we also re-clustered BM and PBMC T and NK cell populations integrated with the overall patient cohort and healthy donors from the HCA dataset **(Fig. 4A)**. Among CD8 T and NK cells, 121,540 BM cells and 147,497 PBMC were analyzed. In the BM, we identified three cytotoxic CD8 T cell subsets (c5, c6, c9) marked by *GZMH* and *IFITM1*, with c9 also showing a type I interferon–responsive signature marked by *TNF* and *IFIT2*, while c6 displayed a CD45RA effector memory phenotype (**Fig. 4B-C, Supplementary Fig 5C, Supplemental table 4-5**). We also identified two lymph-homing subsets (c4, c11; *CCR7*, *NELL2*), a resident-memory like subset (c3; *ZNF683*), a stem-like effector subset (c1; *GZMK*, *RGS1*), and an innate-like subset (c10; *KLRB1*, *IL4I1*).

**Figure 4.**
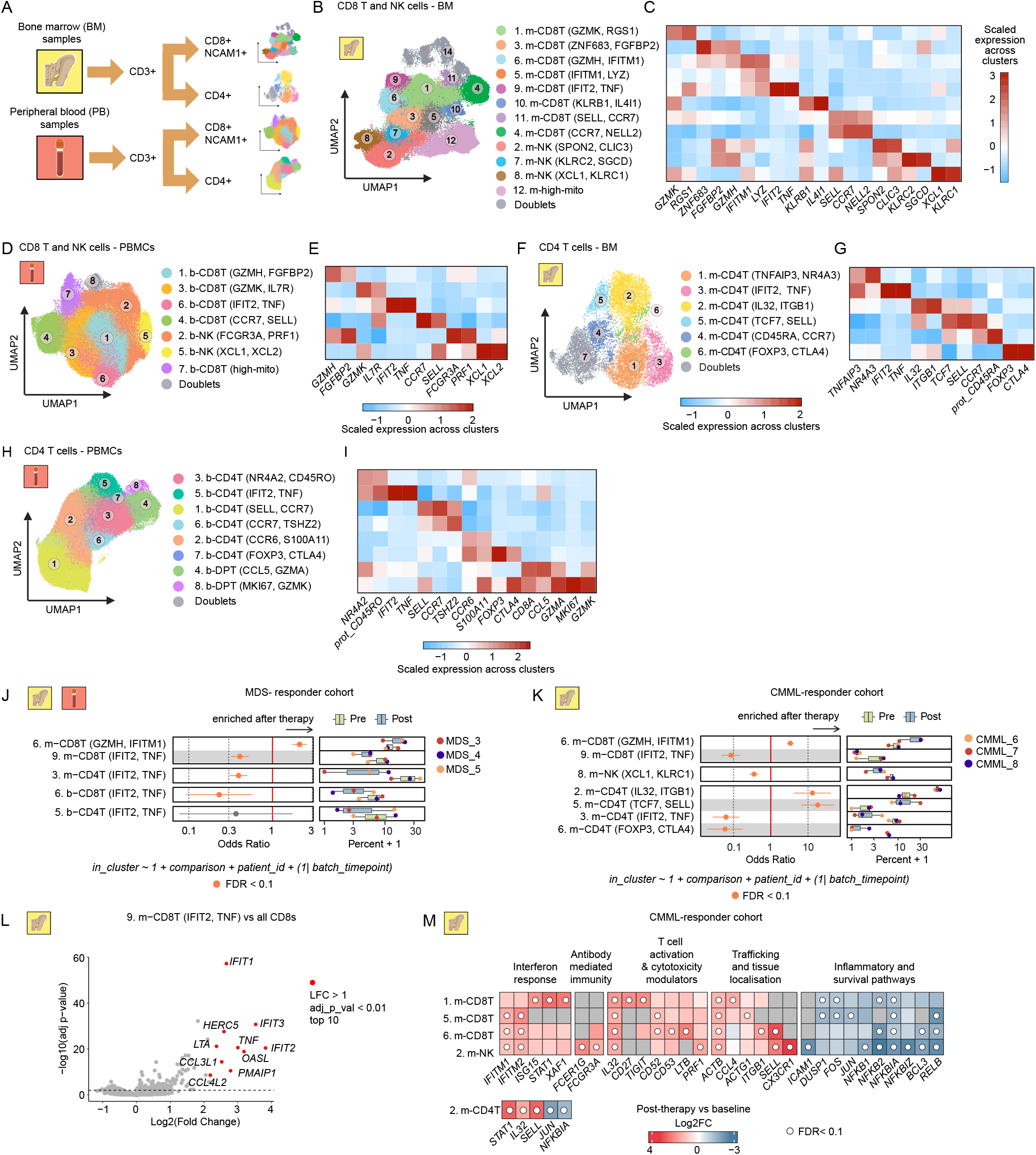
Annotation of CD8 T, NK and CD4 T cell subsets in bone marrow and peripheral blood and their post-therapy abundance and gene expression shifts. **A.** Schematic of the T/NK cell subclustering process. **B.** UMAP embedding of 121,540 CD8 T and NK cells from BM colored by subcluster annotations. **C.** Heatmap showing discriminative markers for each cluster in **B** when compared with every other cluster. Color scale denotes the normalized gene expression (mean zero, unit variance) across cell clusters. **D.** UMAP embedding of 147,497 CD8 T and NK cells from PBMC colored by subcluster annotations. **E.** Heatmap showing discriminative markers for each cell cluster in **D** when compared with every other cluster. **F.** UMAP embedding of 12,627 CD4 T cells from BM colored by subcluster annotations. **G.** Heatmap showing discriminative markers for each cell cluster in **F** when compared with every other cluster. **H.** UMAP embedding of 128,350 CD4 T cells from PBMC colored by annotated subclusters. **I.** Heatmap showing discriminative markers for each cell cluster in **H** when compared with every other cluster. **J-K.** Cell cluster abundance analysis comparing pre- and post-treatment samples for MDS-responders (**J**) and CMML-responders (**K**). Points represent odds-ratio with error bars representing 95% confidence intervals. Color indicates FDR < 0.1 (logistic regression). Box plots show median (line) and IQR, with whiskers no more than 1.5x IQR. **L.** Volcano plot showing the OVA pseudobulk statistics for the m-CD8T (*IFIT2, TNF*) cluster, with the top ten genes by log2FC labeled. **M.** Select DEGs in specific CD8 T and NK clusters when comparing post-treatment to baseline samples in CMML-responders. Color scale indicates log2FC and white dots indicate FDR < 0.1. For **C, E, G and I,** Color scale denotes the normalized gene expression (mean zero, unit variance) across cell clusters.

In PBMCs, we identified a cytotoxic subset (c1; *GZMH*, *FGFBP2*), an effector memory-like subset (c3; *GZMK, IL7R*) and, similar to BM, a type I interferon–responsive subset (c6; *TNF, IFIT2*) along with a lymph-homing subset (c4; *CCR7, SELL*) (**Fig. 4D-E, Supplemental table 6**). Among NK subsets, two conventional subsets were identified in both the BM and PBMC compartments (BM: c7, c8; PBMC: c2, c5), along with a third subset in BM (c2) marked by *SPON2* and *CLIC3*.

Amongst CD4 T cells, sub-clustering of 12,627 BM cells and 128,350 PBMCs identified two lymph-homing subsets in BM (c4, c5), including a naive-like population expressing *CD45RA* and *CCR7* (c4), and a *TCF7, CCR7* central memory-like population (c5) (**Fig. 4F-G, Supplemental table 4-5**). Additional subsets included a stress-responsive population (c1; *TNFAIP3*, *NR4A3*), an effector memory-like subset (c2; *IL32, ITGB1*), and regulatory T cells (Treg; BM: c6; PBMC: c7) marked by *FOXP3*. Similar to CD8 T cells, type I interferon–responsive CD4 T cell subsets were identified in both BM and PBMC (BM: c3; PBMC: c5) characterized by *IFIT2* and *TNF expression*. In PBMCs, additional subsets included a memory-like subset expressing *CD45RO* (c3), two lymph-homing subsets (c1, c6; *SELL, CCR7*), and a *CCR6,S100A11* subset (c2) suggestive of tissue-migratory potential (**Fig. 4H-I, Supplemental table 6**). Among PBMC CD4/CD8 double-positive T cells (DPT), we identified a cytotoxic-like subset (c4; *CCL5*, *GZMA*) and a proliferative subset (c8; *MKI67*). Overall, transcriptional profiles of NK cell, lymph-homing, regulatory T cell, and type I interferon responsive subsets showed substantial overlap between BM and PBMC compartments (OVA pseudobulk FDR < 0.05) (**Supplementary Fig. 5A-B**).

Next, we evaluated treatment-associated changes in cellular abundance and transcriptional states. Amongst responders with MDS, type 1 interferon-responsive and cytotoxic effector memory subsets were enriched at baseline relative to healthy donors (**Supplementary Fig. 5C**). Comparison of pre- versus post-treatment samples demonstrated contraction of *IFIT2, TNF* expressing type 1 interferon-responsive CD8 and CD4 T cell subsets, in both the BM and PBMC compartments following therapy (**Fig. 4J, Supplementary Fig. 5D, Supplemental Table 6)**, along with modest enrichment of the c6-cytotoxic effector memory CD8 T cell subset in BM.

Amongst CMML responders, we observed a similar contraction in type 1 interferon-responsive CD4 and CD8 T cell subsets in the BM post-therapy, together with expansion of the c6-cytotoxic effector memory CD8 T cell subset (**Fig. 4K**). Additionally, BM samples demonstrated reduced Treg abundance and enrichment of naive/central memory (c5) and effector-memory (c2) CD4 T cell subsets (**Fig. 4K, Supplementary Fig. 5D**). The c9 BM CD8 type 1 interferon responsive subset was characterized by elevated expression of interferon-stimulated genes (*IFIT1*, *IFIT2*, *IFIT3*, *OASL*, *HERC5*), pro-inflammatory cytokines (*TNF*, *LTA*), and chemokines involved in immune cell recruitment (*CCL3L1*, *CCL4L2*) (**Fig. 4L, Supplemental Table 4**).

In CMML, amongst responders, combination therapy was associated with upregulated gene expression of interferon response genes (*IFITM1*, *IFITM2*), T cell activation and cytotoxicity modulators (*LTB*), and T cell trafficking (*SELL*) within the c5- and c6-BM cytotoxic CD8 T cell subsets (**Fig. 4M, Supplemental Table 8**). Broad increases in *IL32* expression were observed across multiple T cell subsets, alongside upregulation of *CD27* and *TIGIT* in the stem-like effector T cell subset. NK cells additionally demonstrated increased expression of genes associated with antibody-mediated immune functions, including *FCER1G* and *FCGR3A*. Finally, genes involved in cellular trafficking and tissue localization– including *ACTB*, *CCL4*, *ITGB1*, *SELL*, and *CX3CR1*– were upregulated across selected CD8 and NK BM cell subsets (**Fig. 4M**).

Overall, we identified diverse CD8 and CD4 T cell subsets in this dataset, with often shared transcriptional profiles between PBMC and BM. Following therapy amongst responding patients, we identified a contraction of BM type 1 interferon responsive CD4 and CD8 T cell clusters marked by *TNF/IFIT2* in both MDS and CMML, along with expansion of a cytotoxic effector memory CD8 T cell BM subset.

### Potential predictors of response with combination azacitidine/sabatolimab therapy in MDS

To identify potential baseline predictors of response to combination azacitidine/sabatolimab therapy, we evaluated pre-treatment samples from patients with MDS, comparing those who ultimately responded to therapy (n=3) versus those who did not (n=2) (**Fig. 5A**). Differential abundance analysis revealed baseline differences in CD8 T cells, CD4 T cells, and NK cell subsets (**Supplementary Fig. 6A-B, Supplemental table 6**). Responders had higher proportions of type 1 interferon-responsive CD8 and CD4 T cell subsets in both BM and PBMC compartments, characterized, by *IFIT2* and *TNF* expression, as well as enrichment of a BM CD8 lymph-homing subset with stem-like early memory features **(Fig. 5B).**

**Figure 5.**
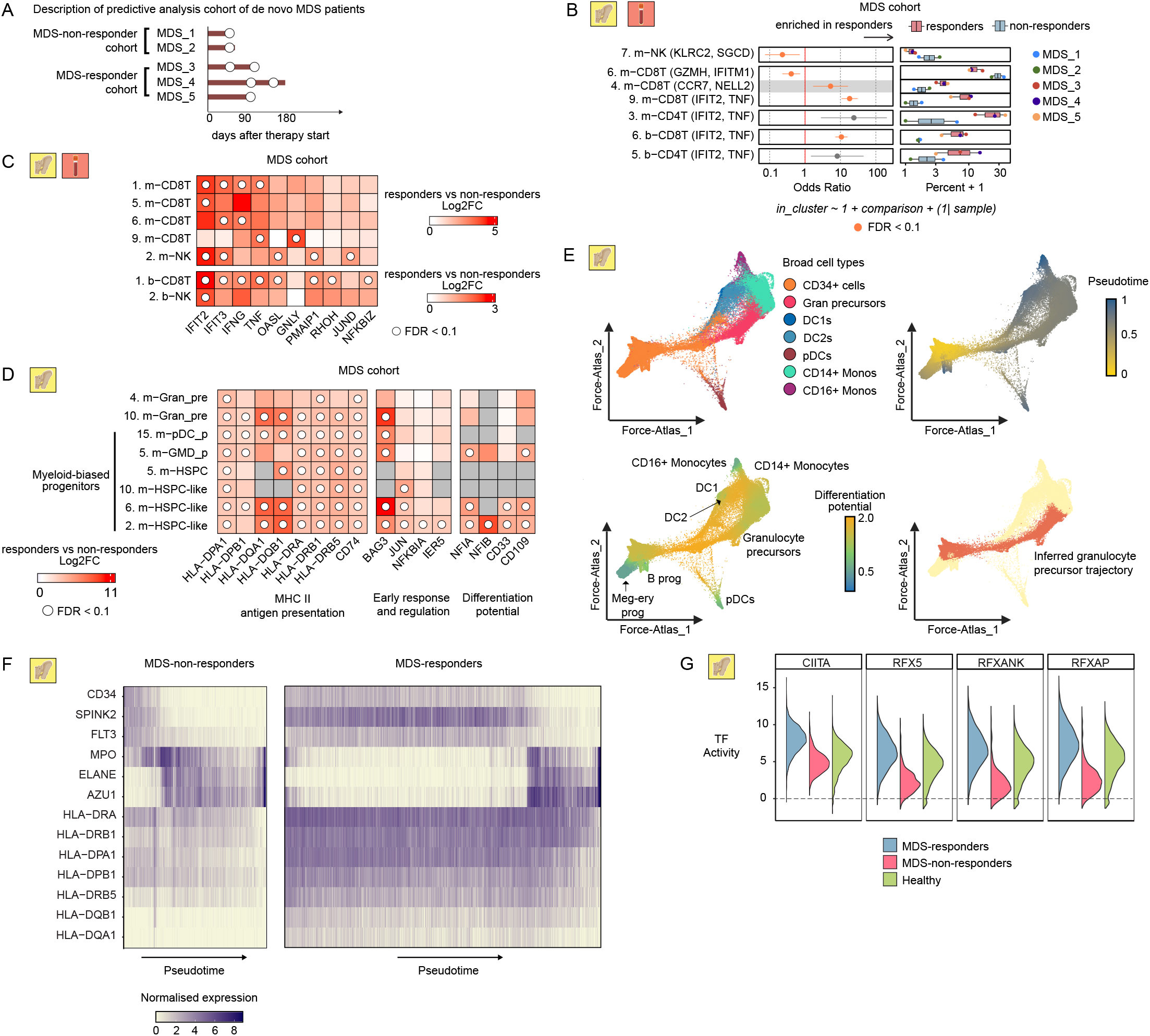
Baseline cellular and transcriptional features distinguishing MDS-responders from non-responders, including MHC class II regulation along the granulocyte trajectory. **A.** Swimmer’s plot showing the de novo MDS cohort used for response-prediction analyses. Baseline pre-treatment samples were used for analyses. **B.** Cell cluster abundance analysis comparing MDS-non-responders and MDS-responders. Points represent odds-ratio with error bars representing 95% confidence intervals. Color indicates FDR < 0.1 (logistic regression). Box plots show median (line) and IQR, with whiskers no more than 1.5x IQR. **C-D.** Select DEGs in specific clusters comparing MDS-responders to MDS-non-responders. Color scale indicates log2FC and white dots indicate FDR < 0.1. **E.** Force-atlas2 embedding of myelomonocytic cells, DCs and HSPCs from BM colored by major cell lineage (top left), inferred pseudotime ordering (top right), defined trajectory end points (bottom left) and inferred granulocyte trajectory (bottom right). **F.** Hexbin plot showing normalized gene expression (Log2CPM) of select markers for hematopoietic stem cells, myeloid-primed MPPs and MHC Class II among granulocyte lineage cells split by response (MDS-non-responders, MDS-responders). Cells (columns) are ordered by inferred pseudotime. **G.** Transcription factor (TF) activity of CIITA, RFXANK, RFXAP, and RFX5 across sample groups at pretreatment timepoint. Half-violin plots display TF activity distributions in the early stages of the granulocyte lineage. Colors indicate groups (MDS-responders, MDS-non-responders or healthy donors), with right-sided violins representing the distribution per group. Data are derived from ULM-estimated TF activities using the CollecTRI network.

At baseline, responders displayed upregulation of immune activation programs across CD8 T cell and NK cell subsets (**Fig. 5C, Supplemental table 8**), including expression of interferon-stimulated genes such as *IFIT2*, *IFIT3*, *OASL*, and *IFNG*. Key effector molecules associated with cytotoxicity and apoptotic signaling, including *TNF*, *GNLY*, and *PMAIP1* were also elevated. In parallel, transcriptional regulators linked to NF-KB and AP-1 signaling – including *JUND*, *NFKBIZ*, and *RHOH* – were upregulated, further supporting the presence of an activated pro-inflammatory immune landscape in responders in these lymphocyte subsets.

Responders also demonstrated increased baseline expression of genes involved in MHC class II antigen presentation across myeloid-primed MPPs, granulocyte-monocyte-DC progenitors, granulocyte precursors, and pDC progenitors in responders, including *HLA-DPA1, HLA-DPB1, HLA-DQA1, HLA-DQB1, HLA-DRA, HLA-DRB1, HLA-DRB5*, and *CD74* (**Fig. 5D, Supplemental table 8**). This was accompanied by increased expression of early response and transcriptional regulatory genes such as *BAG3, JUN, NFKBIA,* and *IER5*, suggesting a primed inflammatory or stress-responsive state within these progenitor populations.

Notably, several HSPC-like clusters (c2, c6, c10) also displayed an upregulation of MHC Class II expression amongst responders at baseline. Increased expression of *NFIA, NFIB, CD33,* and *CD109* was also observed across two HSPC-like subsets and granulocyte-monocyte-DC progenitors. Collectively, these findings suggest activation of antigen presentation and lineage-priming programs within the granulocytic differentiation continuum in responding patients, potentially linked to enhanced immunologic competence at baseline.

To further characterize this differentiation continuum landscape, we generated a force-atlas2 ^43^ embedding trained on diffusion components derived from all BM cells (**Supplementary Fig. 7A**) to visualize inferred developmental trajectories. We subsequently subset CD34 cells, together with monocyte, DCs, and granulocyte precursor branches and recalculated diffusion components to refine lineage resolution (**Fig. 5E**). Diffusion pseudotime analysis identified terminal differentiation states and Palantir^44^ was used to define cells along the granulocyte precursor trajectory (**Fig. 5E**). Mapping normalized expression of key hematopoietic stem cells, multipotent progenitors, granulocyte precursor specification, and MHC class II antigen presentation genes across pseudotime revealed altered differentiation dynamics in non-responders (**Fig. 5F**). Specifically, cells from non-responding MDS patients appeared to bypass the multipotent progenitor and granulocyte precursor MHC-II–high states during progression along the granulocyte lineage.

To further define the regulatory programs underlying these trajectories, we performed transcription factor enrichment analysis in CD34 cells in the early stages of the granulocytic differentiation. Responding patients with MDS demonstrated increased activity of the transcription factor *CIITA*, the master regulator of MHC-II transcription (**Fig. 5G, Supplemental table 9**). In addition, *RFX5*, *RFXANK*, and *RFXAP*, which are core components of the MHC class II transcriptional complex that cooperate with CIITA, were also enriched, supporting activation of coordinated MHC-II transcriptional programs in patients who subsequently responded to therapy.

Thus, collectively these data suggest that upregulation of MHC Class II antigen presentation programs across HSPC, HSPC-like and other subsets across the myeloid differentiation continuum, along with expression of baseline immune activation programs across CD8 T and NK cells, are potential predictors of response to combination therapy in MDS.

## DISCUSSION

We sought to characterize the hematopoietic and immune cell landscape of BM and PBMC compartments in patients with MDS and CMML treated with sabatolimab-based therapy using single-cell RNA sequencing. Integration with publicly available healthy donor datasets enabled refined delineation of hematopoietic and immune subclusters, allowing distinction between populations shared with healthy donors versus those unique to MDS and CMML. Our findings support prior literature of baseline differences in acquired immune subsets (B and T cells) as well as innate immune subsets (NK cells, CD16 monocytes) in MDS and CMML,^45,46,47^ along with findings of left shifted impaired hematopoiesis, with CD34 enrichment, along with a myelomonocytic differentiation bias. While the overall HSPC-like compartment exhibited a myelomonocytic differentiation bias, individual subclusters retained transcriptional features resembling stem cells, MPPs, or MEPs, similar to trajectory biases reported in CMML HSPCs.^44,48^

The pathogenic role of chronic inflammation has been recognised in MDS pathogenesis.^49^ Integration of scRNAseq data with healthy donor datasets helped to distinguish a disease-associated CD34 HSPC-like population in MDS and CMML characterized by AP-1 transcription factor activity, NF-κB–associated inflammatory signaling and features suggestive of impaired differentiation. Enrichment of NF-κB and immediate early-response genes supports the presence of a stress-adapted progenitor state in myeloid malignancy. We also observed context-specific remodeling following therapy. In MDS, the HSPC-like population retained stress-response and metabolic adaptation programs post-treatment, whereas in CMML we observed downregulation of inflammatory and stress-response pathways within dendritic cells, dendritic cell progenitors, and antigen-presenting monocyte populations amongst responders, suggesting that therapy may attenuate NF-κB/AP-1–driven inflammatory myeloid program in CMML.^50–52^ The difference in median post-treatment collection timeframes between MDS and CMML may have impacted these findings. Further, responders demonstrated increased CD16 monocytes in CMML and expansion of erythroid progenitors and erythroid cells in MDS following therapy, suggesting partial restoration of monocytic and erythroid differentiation programs. Overall, these data support clinical investigation of NF-κB modulation, particularly in CMML, and it will be important to assess if such modulation primarily improves cytopenias or has the potential for disease course modification. While we could not assess direct mutational assignment to specific subclusters such as those within the HSPC-like compartment due to sequencing depth, CNV analyses confirmed multilineage neoplastic clonal hematopoiesis with myeloid predominance.^53,54^

Relatively little is known regarding the immunologic effects of TIM-3 inhibition combined with hypomethylating agents in patients with myeloid malignancies. Our data support reported findings of TIM-3 expression on DCs and NK cells, with additional expression across CD34 cells, monocytes, and activated T cell populations, supportive of a key connection between APC and innate immunity with TIM-3 biology. Huuhtanen *et al.* evaluated decitabine with sabatolimab primarily in relapsed/refractory AML and found similar observations of TIM-3 expression across NK, dendritic, myeloid, and activated T cell populations.^55^ That study also reported post-treatment dendritic cell expansion and inflammatory pathway activation in responders.^55^

In the present study, a key observation was the association between elevated MHC-II expression across several populations including HSPCs and HSPC-like cells and enrichment of MHC-II transcriptional regulators with therapeutic response to combination azacitidine/sabatolimab therapy in MDS. These findings suggest that antigen presentation outside of TIM-3 expression itself may play a role in response. Although MHC-II expression on HSPCs and leukemia cells has been described,^56–58^ its functional significance has only recently begun to emerge. Hernandez-Malmierca *et al.* demonstrated that HSPCs are capable of activating antigen-specific CD4 T cells through MHC-II–dependent antigen presentation, supporting a potential immunosurveillance role for progenitor populations.^59^ Our trajectory analyses further showed that non-responders bypassed MHC-II–high developmental states along the granulocytic continuum, suggesting impaired maturation and potentially reduced antigen presentation capacity. Due to limited quantities of longitudinal post-therapy samples, we could not functionally test how MHC-II expression was associated with response, such as testing of progenitor or DC antigen presentation interactions with activated T cells. Since hypomethylating agents can modulate HLA expression and antigen presentation,^60–62^ these findings provide rationale for further investigating the role of MHC-II in maturation itself and combination strategies that enhance MHC-II expression or antigen presentation to improve responses to therapy.

Another notable finding was the identification of a type I interferon–responsive T cell population in both BM and PBMC compartments and across CD4 and CD8 T cell subsets. Among patients with MDS, this population was enriched at baseline in eventual responders relative to non-responders, potentially reflecting a pre-existing inflammatory effector state or primed adaptive immune response. Similar predictive inflammatory T cell populations have been described in solid tumors receiving immunotherapy.^63–65^ Interestingly, this population contracted following therapy in both MDS and CMML, potentially reflecting an attenuation of aberrant inflammatory signaling which may facilitate improvement of cytopenias.^66^ Since these cells had a specific gene signature, it would be of interest to see if these populations and/or enriched MHC-II pathway expression could be prospectively identified in order to help identify and select patients who may be more likely to benefit to accrue to studies evaluating HMA/TIM-3 inhibition, as other TIM-3 modulating therapies are being explored.

Responders also demonstrated evidence of enhanced NK-cell cytotoxicity and increased interferon-responsive, trafficking-associated, and cytotoxic gene expression programs across several CD8 T cell subsets following therapy, suggesting enhanced effector activity. The CD4 T cell compartment demonstrated reduced Treg abundance alongside expansion of *IL32* helper/effector populations and *TCF7*-expressing subsets, which have been associated with immune checkpoint blockade responses.^67^ Collectively, these findings support enhanced adaptive and innate immune activation in responding patients.

This study has several limitations. Samples were retrospectively obtained from an institutional tissue bank, and although pretreatment BM samples were prioritized, available post-treatment timepoints varied across patients in this exploratory cohort. CITE-seq profiling was feasible only in a limited subset due to sample constraints, preventing comprehensive paired protein analyses. Response was defined as stable disease or better, which may encompass biologically heterogeneous outcomes. However, treatment-associated changes were observed across responding patients, suggesting shared underlying biology. Given most patients received combination therapy, we were unable to fully disentangle the independent contributions of sabatolimab and hypomethylating therapy. Although limited samples from patients receiving sabatolimab monotherapy or historical HMA therapy alone were included for cluster definition purposes, sample sizes were insufficient for meaningful comparative analyses. Additionally, because we tried to include longitudinal samples, our overall combination therapy cohort consisted of 6 subjects who had response to therapy and 2 that did not, so our results may be biased towards findings observed in responders. While the findings of similar OS in the randomized phase III study comparing azacitidine plus sabatolimab with azacitidine plus placebo (NCT04266301)^37^ suggests that additional components beyond immune factors contributed towards lack of response or loss of response, the baseline features that were found to be associated with response provide avenues for future therapeutic exploration.

In summary, we identified widespread immune and hematopoietic remodeling in patients with MDS and CMML treated with azacitidine and sabatolimab. Baseline disease states were characterized by reduced adaptive immune cell populations, impaired myeloid differentiation, and inflammatory transcriptional programs across monocyte, dendritic, and HSPC-like compartments. Responders demonstrated evidence of restored differentiation, enhanced innate and adaptive immune activation, and increased baseline expression of MHC-II programs. Together, these findings support an important role for coordinated interactions between myeloid progenitors, innate immunity, and adaptive immune responses in myeloid malignancies. These findings suggest that inflammatory progenitor states and immune–hematopoietic interactions represent biologically relevant therapeutic axes in MDS and CMML that warrant further investigation.

## METHODS

### Study Samples

Institutional review board approval (MGB Insight #2020P000656) was obtained for retrospective chart review, data collection and secondary biospecimen use from the MGH hematologic malignancies tissue bank of samples that had been previously collected with informed consent (DFHCC #13-583) from subjects with MDS and CMML who had been treated with either hypomethylating (HMA) therapy alone as part of standard care or from subjects treated at MGH who had received investigational therapy on study NCT03066648. Briefly, samples had been collected in EDTA followed by buffy coat extraction after centrifugation. After application of red blood cell lysis buffer, cells were centrifuged, pelleted, and resuspended either in a subject’s own plasma or fetal bovine serum combined with DMSO (final concentration 7.5%) followed by cryofreezing for long term use.

NCT03066648 was a phase Ib, multi-center, open label, and multi-arm study evaluating the safety and tolerability of sabatolimab as monotherapy, sabatolimab in combination with spartalizumab (PDR001, an anti-PD1 inhibitor), or sabatolimab combined with HMA therapy (either azacitidine or decitabine) for patients with AML, HR- or vHR-MDS, and CMML^35^. Additional treatment arms and full dosing details have been previously described, but briefly patients on the monotherapy arm received sabatolimab intravenously (IV) 400 mg or 1200 mg every 2 weeks, and patients on the sabatolimab plus HMA combination therapy arm were treated with escalating doses of sabatolimab at either 240 mg or 400 mg every 2 weeks or 800 mg every 4 weeks IV with either azacitidine (75 mg/m2/day, IV or subcutaneous on days 1-7) or decitabine (20 mg/m2/day, IV on days 1-5) in 28-day cycles.

### Sample selection and characteristics

For this study, scRNA-seq and cellular indexing of transcriptomes and epitomes by sequencing (CITE-seq) data were generated from viably cryofrozen BM and PBMC samples from 16 patients (**Fig. 1A**). Samples were primarily selected based on pretreatment baseline sample availability. 12 patients had been previously treated on NCT03066648 including 4 patients with RR-MDS treated with either sabatolimab alone (n=3) or combined with spartalizumab (n=1), and 8 patients treated with azacitidine combined with sabatolimab (n=5 de novo MDS, n=3 CMML). Samples included pretreatment baseline samples and those collected either during early treatment timepoints (typically from the end of cycle 1 (C1) or C2), later treatment time points (typically from the end of C4-C6), or at the end of therapy (EOT) (**Supplementary Tables 1-2**). Samples from 4 patients with either de novo MDS (n=2) or CMML (n=2) treated with HMA therapy alone (either azacitidine or decitabine) as part of standard care were also evaluated (“historical HMA-alone” samples), primarily with pretreatment baseline BM samples being available (**Supplementary Tables 1-2**). Retrospective chart review was used to ascertain best overall response to therapy using International Working Group (IWG) MDS 2006 criteria^68^. Best responses of stable disease (SD) or better were characterized as responders for the purposes of this study (**Supplementary Table 2**). Clinical karyotype evaluation and next generation sequencing (NGS) molecular mutation data were retrospectively notated from the time of diagnosis or pretreatment samples (**Fig 1B**).

### PBMC and bone marrow specimen processing and CITE-seq staining

Cryopreserved PBMC and BM samples were thawed at 37°C, diluted with a 10x volume of RPMI with 10% heat-inactivated human AB serum (Sigma Aldrich), and centrifuged at 300 g for 7 mins. To improve recovery of viable cells and remove debris, we performed density gradient centrifugation with Ficoll-Paque (Fisher Scientific) for every specimen. The cellular fraction was washed by centrifugation and cells were resuspended in CITE-seq buffer (RPMI with 2.5% [v/v] human AB serum and 2 mM EDTA) and added to 96-well plates. Additional dead cells were removed with an Annexin-V-conjugated bead kit (Stemcell Technologies, 17899) and red blood cells were removed with a glycophorin A-based antibody kit (Stemcell Technologies, 01738); modifications were made to the manufacturer’s protocols for each to accommodate a sample volume of 150 μL. Cells were quantified with an automated cell counter (Bio-Rad, TC20), after which, 2.5 x 10^5 cells were resuspended in CITE-seq buffer containing TruStain FcX blocker (BioLegend, 422302) and MojoSort CD45 Nanobeads (BioLegend, 480030). Cells were then washed three times with CITE-seq buffer using a magnet to retain cells. Live cells were counted with trypan blue, and samples from each cohort bearing different hashtag antibodies were pooled together at equal concentrations. All samples were pooled in groups of eight and loaded on one 10x Genomics channel; all 8 samples for each pool were balanced for disease status, sex, treatment conditions, treatment response outcome and treatment time point. Leftover cells were lysed in Buffer TCL + 1% beta-mercaptoethanol (BME) at a concentration of 1,000 cells/uL to enable bulk RNA-sequencing (see genetic demultiplexing section below). Note that the PBMC specimens were pooled together, and the BM samples were pooled together. Pooled samples were filtered with 40 μM strainers, centrifuged, and resuspended in CITE-seq buffer with TotalSeq-C antibody cocktail (BioLegend; **Table S10**). Cells were incubated on ice for 30 mins, followed by three washes with CITE-seq buffer and a final wash in the same buffer without EDTA (RPMI with 2.5% [v/v] human AB serum). Cells were resuspended in this buffer without EDTA, filtered a second time, and counted.

### Single-cell gene expression, feature barcode, and sequencing

An input of 50,000 hashed PBMC or BM samples were loaded on a single channel in the 10X Chromium instrument for blood immune cell analysis, aiming for a recovery goal of 25,000 single cells. Single-cell libraries were generated with the Chromium Single Cell 5’ kit (V2 NextGEM 10X Genomics, PN-1000263) together with the 5’ Feature Barcode library kit (10X Genomics PN-1000256) according to manufacturer’s protocols. The library quality was assessed with an Agilent 2100 Bioanalyzer. All gene expression and feature barcode were sequenced on an Illumina NovaSeq using the S4 300 cycles flow with the following sequencing parameters: (read 1=26; read 2 = 90; index 1 =10; index 2 =10). Measurements from the two modalities were measured in the same cells and could be associated using the cell barcode.

### Genetic demultiplexing of pooled data

An aliquot of cells from each PBMC and bone marrow specimen was lysed in Buffer TCL + 1% beta-mercaptoethanol (BME) at a concentration of 1,000 cells/uL. We used the leftover cells from each specimen that were not pooled together in groups of eight and loaded on one 10x Genomics channel. 25 uL of lysed cells per sample was then used to generate low-input bulk RNAseq using the SmartSeq2 protocol ^69^ to enable genetic demultiplexing. Bulk RNAseq data was sequenced on a NextSeq 550 with the parameters: read 1=38; read 2=38; index 1=8; index 2=8. Note that only 1 sample per donor was profiled for downstream demultiplexing (i.e. 1 time point for either bone marrow or PBMC specimen per donor). To assign cells from pooled samples back to individual donors, we first applied Souporcell ^70^ to the raw aligned BAM files to perform unsupervised clustering based on common genetic variants (SNPs). Souporcell identifies genotype clusters by modeling allele fractions in each cell, allowing for robust clustering even in the presence of ambient RNA (soup). The resulting clusters represent putative donor groups but are unlabeled with respect to individual identities. To resolve the identity of each cluster, we leveraged donor genotype information. Using Smart-seq2 bulk-RNA-seq runs, we used cellSNP-lite (v1.2.0) to extract SNP-level allele counts for each donor in the pooled samples. These individual-specific SNP profiles were used as reference genotypes. We then applied VireoSNP^71^ to map Souporcell clusters to individual patients, using a supervised approach that assigns cells to individuals by matching their observed SNPs to the reference profiles. This allowed us to annotate each Souporcell cluster with its corresponding patient identity with high confidence. Cells that did not clearly map to a single genotype or had high levels of ambient RNA were excluded from downstream analysis.

### scRNA-seq read alignment and quantification

Raw sequencing data, both from samples sequenced in-house and public data from healthy donors, was pre-processed with CellRanger (6.0.1, 10x Genomics) to demultiplex FASTQ reads, align reads to the human reference genome (GRCh38, v3.0.0 from 10x Genomics), and count unique molecular identifiers (UMI) to produce a cell x gene count matrix ^72^. All count matrices were then aggregated with Pegasus (v1.1.0, Python) using the *aggregate_matrices* function ^73^. Low-quality droplets were filtered out of the matrix prior to proceeding with downstream analyses using the percent of mitochondrial UMIs and number of unique genes detected as filters (20% mitochondrial UMIs, > 300 unique genes). The percent of mitochondrial UMI was computed using 13 mitochondrial genes (MT-ND6, MT-CO2, MT-CYB, MT-ND2, MT-ND5, MT-CO1, MT-ND3, MT-ND4, MT-ND1, MT-ATP6, MT-CO3, MT-ND4L, MT-ATP8) using the *qc_metrics* function in Pegasus. The counts for each remaining cell in the matrix were then log-normalized by computing the log1p(counts per 100,000), which we refer to in the text and figures as logCPM. The detailed quality control statistics for these datasets are compiled in **Supplementary Table 1**.

### Integration of scRNA-seq data from the Human cell atlas

Raw scRNA-seq data from adult human peripheral blood and bone marrow samples were downloaded from the Human Cell Atlas project ^39^. These files were preprocessed following the identical pipeline described previously. To balance class sizes between healthy and myeloid malignancy samples, the blood dataset was randomly subsampled to 200,000 cells prior to downstream analysis. Harmony^74^ was used to correct the principal components for study-related batch effects.

### Basic clustering

2,000 highly variable genes were selected using the highly_variable_features function in Pegasus and used as input for principal component analysis. We ran the molecular cross-validation (MCV) algorithm^75^ to select the number of components for downstream analysis. To account for technical variability between donors, the resulting principal component scores were aligned using the Harmony algorithm^74^. The resulting principal components were used as input for Leiden clustering and UMAP algorithm (spread = 1, min-dist = 0.5).

### Marker gene identification and cell annotation

The marker genes defining each distinct cell subset from our global and lineage-specific subclustering analyses were determined by applying two complementary methods. First, we calculated the area under the receiver operating characteristic (AUROC) curve for the logCPM values of each gene as a predictor of cluster membership using the *de_analysis* function in Pegasus. Genes with an AUROC ≥ 0.75 were considered marker genes for a particular cell subset. Second, we created a pseudobulk count matrix^76^ by summing the UMI counts across cells for each unique cluster/sample combination, creating a matrix of n genes x (n samples*n clusters). We performed “one-versus-all” (OVA) differential expression (DE) analyses for each cell subset using the Limma package (v3.54.0, R)^77^. For each subset, we used an input model *gene ∼ in_clust*, where *in_clust* is a factor with two levels indicating if the sample was in or not in the subset being tested. A moderated t-test was used to calculate P values and compute a false discovery rate (FDR) using the Bejamini-Hochberg method. We identified marker genes that were significantly associated with a particular subset as having an FDR<0.01 and a log2 fold change > 1. The AUROC and OVA pseudobulk marker genes for all cell subsets can be found in Supplementary tables 4-5.

### Abundance analysis

We used a mixed-effects association logistic regression model similar to that described by Fonseka et al^78^ to identify cell abundance associations comparing 1) malignancy (MDS/CMML) versus healthy donor status (comparing baseline samples), 2) predictiveness of response to therapy (comparing baseline samples from eventual responders versus non-responders to therapy) and 3) timepoint associations (comparing pre-treatment baseline samples versus post-treatment samples). We used the *glmer* function from the lme4 package (v1.1-31, R) to fit a logistic regression model for each cell subset. Each subset was modeled independently with a “full” model as follows:

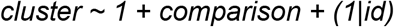

where *cluster* is a binary indicator set to 1 when a cell belongs to the given cell subset or 0 otherwise, *comparison* is a factor with 2 levels which represented either the malignancy (MDS/CMML versus healthy donor at baseline), response predictiveness (responders versus non-responders) or timepoint (pre-treatment versus post-treatment), and *id* is a factor indicating the donor or sample. The notation *(1|id)* indicates that *id* is a random intercept. To determine significant associations, a “null” model of cluster ∼ 1 + (1|id) was fit and a likelihood ratio test was used to compare the full and null models. A false discovery rate was calculated using the Benjamini-Hochberg approach and clusters with an FDR<10% were considered significant. All abundance analysis results can be found in **Supplementary table 6**.

### Differential gene expression analysis

Differential gene expression comparisons of baseline samples between eventual responders versus non-responders, or pre-treatment versus post-treatment were performed on pseudobulk count matrices using the DESeq2 package (v. 1.38.3, R) ^79^. The input model was *gene ∼ comparison*, where *comparison* was a binary variable indicating which group (i.e. responder/non-responder, pre-treatment/post-treatment) the sample belonged to. Significantly differentially expressed genes (DEGs) were identified using a Wald test (FDR < 0.1). DEGs were visualized with the ComplexHeatmap package (v.2.15.4, R). All DEG results can be found in **Supplementary table 8**.

### Trajectory inference with ForceAtlas2, diffusion maps, and Palantir

To investigate differentiation trajectories in myeloid lineages, we first performed trajectory analysis on all BM cells to identify relevant end points. A shared nearest neighbor (SNN) graph was constructed on the Harmony-corrected PCA components. For manifold learning, we applied ForceAtlas2 (FA) ^43^ layout embedding to capture global transcriptional relationships, revealing transcriptional movements from CD34 cells toward discrete lineage-specified endpoints, including monocyte–granulocyte-DCs, erythroid cells, megakaryocytes, and lymphoid cells.

To further identify branching trajectories within the monocyte–granulocyte-DC endpoint, we repeated these steps with the CD34 cells, monocytes, granulocyte precursors and DCs. To infer pseudotime and lineage probabilities, we used the Palantir ^44^ framework. Diffusion maps were computed with 20 components, and multiscale diffusion distances were estimated using Palantir’s built-in tools. We defined the trajectory starting point as CD34 hematopoietic progenitor cells with median coordinates in the FA embedding. Terminal states were selected based on representative cells from granulocyte precursors, classical and non-classical monocytes, cDC1, cDC2, and pDCs. Palantir was run using 1200 waypoints and a kNN value of 100 to compute pseudotime, entropy, and fate probabilities across inferred terminal lineages. Trajectory branches were visualized on the FA embedding, and branch-specific gene expression dynamics were computed using spline-based trend fitting. Trajectories were defined as cells with a specific trajectory probability greater than 0.25.

### Inference of gene expression programs with batch-corrected cNMF

We applied batch-corrected consensus non-negative matrix factorization (cNMF) ^80^ to the myelomonocytic and DC subset to identify shared gene expression programs (GEPs) across cells. Batch correction was performed during preprocessing using the built-in Harmony integration in the cNMF pipeline to account for donor- and sample-level variation. The number of programs (K) was selected based on the tradeoff between solution stability and reconstruction error across multiple NMF replicates. Following cNMF decomposition, we filtered low-quality or redundant factors using a Euclidean distance threshold of 0.18 between gene loadings across factors to collapse near-duplicate programs. Consensus programs were then generated by aggregating stable and distinct components across replicates. For interpretability, we computed the mean usage of each GEP across annotated cell types and visualized these distributions to identify GEPs associated with patient-specific monocyte subsets. Top 50 genes from each consensus gene expression program can be found in **Supplementary Table 7.**

### Transcription factor activity analysis using CollecTRI

Transcription factor (TF) activity inference was performed using the CollecTRI ^81^ regulatory network and Univariate Linear Model (ULM) approach. The human CollecTRI network was obtained with transcription factor complexes maintained as single regulatory units, then preprocessed by removing duplicate source-target pairs and entries with missing values. TF activities were estimated using *run_ulm()* from the decoupler package ^82^, which calculates activity scores through linear regression between normalized gene expression levels and regulatory network weights for each TF-target relationship. Analysis focused on early hematopoietic cells in the granulocyte precursor lineage. Cells were stratified by pretreatment status, with cells from healthy baseline donors included as reference populations.

Differential TF activity analysis was performed using decoupler’s *rank_sources_groups()* with t-test corrected for overestimated variance, comparing each treatment group against all others combined. The top 20 transcription factors with significant activity changes were selected per group. Results were visualized using violin plots displaying selected TF activity score distributions across treatment groups.

### Copy number variation analysis using NUMBAT

We inferred chromosomal abnormalities at single-cell resolution using NUMBAT^83^, a probabilistic tool for detecting subclonal copy number alterations (CNAs) in scRNA-seq data. An initial unbiased analysis identified CNA profiles in subclonal populations that showed strong concordance with clinical cytogenetics results in some samples. To enhance sensitivity, we then re-ran NUMBAT on all samples using fixed chromosomal boundaries and copy number states derived from a predefined clinical cytogenetics panel. This guided approach recovered a greater number of CNA-positive cells across patients and timepoints, improving detection of low-prevalence subclones and providing a more detailed view of clonal architecture.

## DATA AVAILABILITY

scRNA-seq count matrices from this study and related data is deposited in the GEO database (under accession GSE343597), and raw human sequencing data will be available in the controlled access repository dbGaP (https://www.ncbi.nlm.nih.gov/gap) upon publication of this manuscript.

## CODE AVAILABILITY

Source code for data analysis will be available on GitHub upon publication of this study at https://github.com/villani-lab/anti_tim3. A full list of software packages and versions included in the analyses is included in **Supplementary Table 11**.

## AUTHOR CONTRIBUTIONS

R.N., A.M.B., and A.C.V. conceived the study. R.N. and A.C.V. led the overall study design; A.D. led all analyses with contributions from N.S., T.C., S.A.F; A.T. and B.A. performed all experimental work; cell annotation review was performed by A.D., N.S., R.N., and A.C.V.; J.L.S. and T.C. assisted with tissue bank sample processing; J.L.S., T.T.B., and R.N. completed retrospective chart review; A.T.F., G.S.H., H.H., P.C.A., A.M.B., T.A.G., and R.N., assisted with the sample cohort, clinical expertise and/or clinical research protocols; A.D., N.S., R.N., and A.C.V. led data interpretation with assistance from T.S., P.R., M.J,. L.B., and A.M.B.; R.N. and A.C.V. managed and supervised the entire study; A.D., N.S., R.N., A.C.V. wrote the manuscript with input from all authors.

## ACKNOWLEDGEMENTS

We thank Gary Reynolds, Steven Blum, Rachelly Normand for valuable discussions on the project. We thank Kamil Slowikowski, Gary Reynolds for their technical support on this project. We thank valuable assistance from clinical research assistants, trial coordinators, and nurses from the MGH Center for Leukemia. This work was made possible by the generous research support from the Novartis Institutes for BioMedical Research (R.N. and A.C.V.), the National Institute of Health Director’s New Innovator Award (DP2CA247831; to A.C.V.), the Massachusetts General Hospital Transformative Scholar in Medicine Award (to A.C.V.), and the MGH Howard M. Goodman Fellowship (to A.C.V.).

## DECLARATION OF INTERESTS

A.C.V. has received an honorarium from Bristol Myers Squibb; and financial interest in 10X Genomics. 10X Genomics designs and manufactures gene sequencing technology for use in research, and such technology is being used in this research; these interests were reviewed by The Massachusetts General Hospital and Mass General Brigham in accordance with their institutional policies. R.N. reports research funding to the institution from Novartis, Janssen, BMS, and Cullinan Florentine, and spouse employment with Sanofi Genzyme. A.T.F reports consulting activity for Novartis, Servier, BMS, Pfizer, Genmab, Autolus, Daiichi Sankyo, Astellas, Johnson & Johnson, Abbvie, Genentech, AstraZeneca, Kura, ThermoFisher, Azurity, BeOne, Syndax, Ipsen; and clinical trial support from Servier, BMS, Kura, Abbvie, Genentech. T.A.G reports spouse employment with AstraZeneca. G.S.H reports consulting work for Novartis, BMS, GSK, Sobi, Incyte, Damora, Calytryx, Cogent, Merck, DISC, Takeda and Pharmaessentia. A.M.B. reports consulting for Novartis, Servier, BMS, Geron, Syndax, Takeda, Taiho, Sanofi, Alexion, Abbvie, Keros, Rigel. The remaining authors declare no competing interests.

## Tables

**Table S1. Sample information for bone marrow and peripheral blood single-cell data.**

Sample-level metadata. Includes unique sample identifier (Sample), corresponding subject (Subject ID), sequencing batch (Batch), timepoint of collection (Timepoint), tissue of origin (Tissue), number of post-QC cells (Number of cells post-qc), whether the sample was processed as a single subject or pooled channel (Single or pooled channel), and whether CITE-Seq data was generated (CITE-Seq).

**Table S2. Subject demographics and disease characteristics, along with sample collection timepoints and disease status per timepoint**.

Clinical demographic metadata for each subject at baseline including subject identifier (Subject ID), demographic information (Age, Sex), disease diagnosis (Disease), and treatment assignment (Treatment). Also noted is an overview of sampling strategy and disease assessments across study, with each row representing a sample timepoint per subject, including overall response assessment (Overall Response), best response achieved (Best Response), numerical labels for clinical assessment timepoints in days (Disease Assessment Timepoints (days)), corresponding abbreviated labels (Timepoint abbreviation), response at the timepoint (Response at Timepoint), and whether Bone Marrow or Peripheral Blood was collected. Related to Fig 1.

**Table S3. Library information for bone marrow and peripheral blood single-cell data**

Information about single-cell libraries used for gene expression and CITE-seq profiling. Each row includes the sequencing batch identifier (Batch), gene expression index (GEX_Index), and feature barcode index (FBC_Index). Related to Fig 1.

**Table S4. Full marker genes**

Each row represents a gene used to assign a granular cell type to a given cluster or to identify a broad lineage. Marker gene statistics include AUROC, log2 fold-change (logFC), raw p-value (p_val), and FDR-adjusted p value (padj). Cell types are nested under broader categories used for downstream grouping and tissue. Related to Figs 2, 3.

**Table S5: Curated genes used for annotations including references**

Manual annotation of single-cell clusters across tissues and embeddings. Each row corresponds to a cluster, with information on tissue of origin (Tissue), embedding type or analysis context (*Embedding*), broad lineage classification (*Broad annotation)*, numeric cluster identifier (Cluster name), an alternate label (Synonym), canonical markers for lineage (Lineage markers), shared gene used to annotate multiple clusters (Multi-cluster specific markers), specific markers used to refine cluster annotation (Cluster specific markers), Top marker genes using pseudobulk_log2FC>1, padj<0.1 and auroc >0.5, supporting references (Relevant references) and additional descriptions. Related to Figs 2, 3, 4

**Table S6: Abundance analysis results.**

Statistical results from cell-type abundance analyses across multiple comparisons. Each row represents a unique combination of cell type/major lineage and comparison (e.g., Pre vs Post MDS, Pre vs Post CMML, CMML baseline vs Healthy) with corresponding effect sizes and significance testing. Reported metrics include the odds ratio (OR), 95% confidence interval bounds (confint_95_lower, confint_95_upper), p-value (pval), and FDR-adjusted p-value (padj). Metadata includes major lineage classification (major lineage), comparison label (comparison), embedding level (embedding_level), and tissue compartment (tissue). Related to Figs 1, 3, 4, 5.

**Table S7. Top 50 genes from consensus gene expression programs (GEPs).**

Ranked list of top 50 genes by gene weights for each gene expression program. Weights are calculated as the median gene loading across multiple NMF runs and L1-normalized within each program. Related to Fig 2.

**Table S8. Differential gene expression results.**

Differential gene expression results across comparisons and cell types. Each row corresponds to a unique gene within a given cell type and comparison (e.g., CMML Responders vs Healthy, MDS Responders vs Non-Responders). Reported metrics include the average normalized expression (baseMean), effect size (log2FoldChange), standard error (lfcSE), test statistic (stat), raw p-value (pvalue), and FDR-adjusted p-value (padj). Metadata columns specify the comparison label (comp), embedding of cell type (embedding), and tissue compartment (tissue). Related to Figs 2, 3, 4, 5.

**Table S9. Top transcription factors enriched in each group.**

Results of transcription factor enrichment analyses for each group (e.g., MDS-Res vs all, MDS-NonRes vs all, Healthy vs all). Each row represents a transcription factor (TF) identified as significantly enriched. Reported statistics include the enrichment score (Stat), raw p-value (P_value), effect size as mean change (Mean_change), and FDR-adjusted significance (Adj_p_val). Metadata specifies the comparison label (comparison). Related to Fig 5.

**Table S10. Antibody panel metadata for CITE-seq profiling.**

Each entry lists the unique DNA oligonucleotide identifier assigned to the antibody (Cite-seq DNA ID), the manufacturer-provided antibody name and species reactivity (Antibody description), the antibody clone designation (Clone), the DNA barcode sequence conjugated to the antibody (Barcode), the target surface molecule (Protein Target), and the corresponding approved HUGO Gene Nomenclature Committee symbol (Approved Gene symbol). Related to Fig 1.

**Table S11. Software.**

List of all python and R libraries used for analyses and plotting. Related to all Figs.

