## Supplementary Figures for "Response to azacitidine and the TIM-3 inhibitor, sabatolimab, in MDS and CMML is associated with distinct immune and antigen-presentation states"

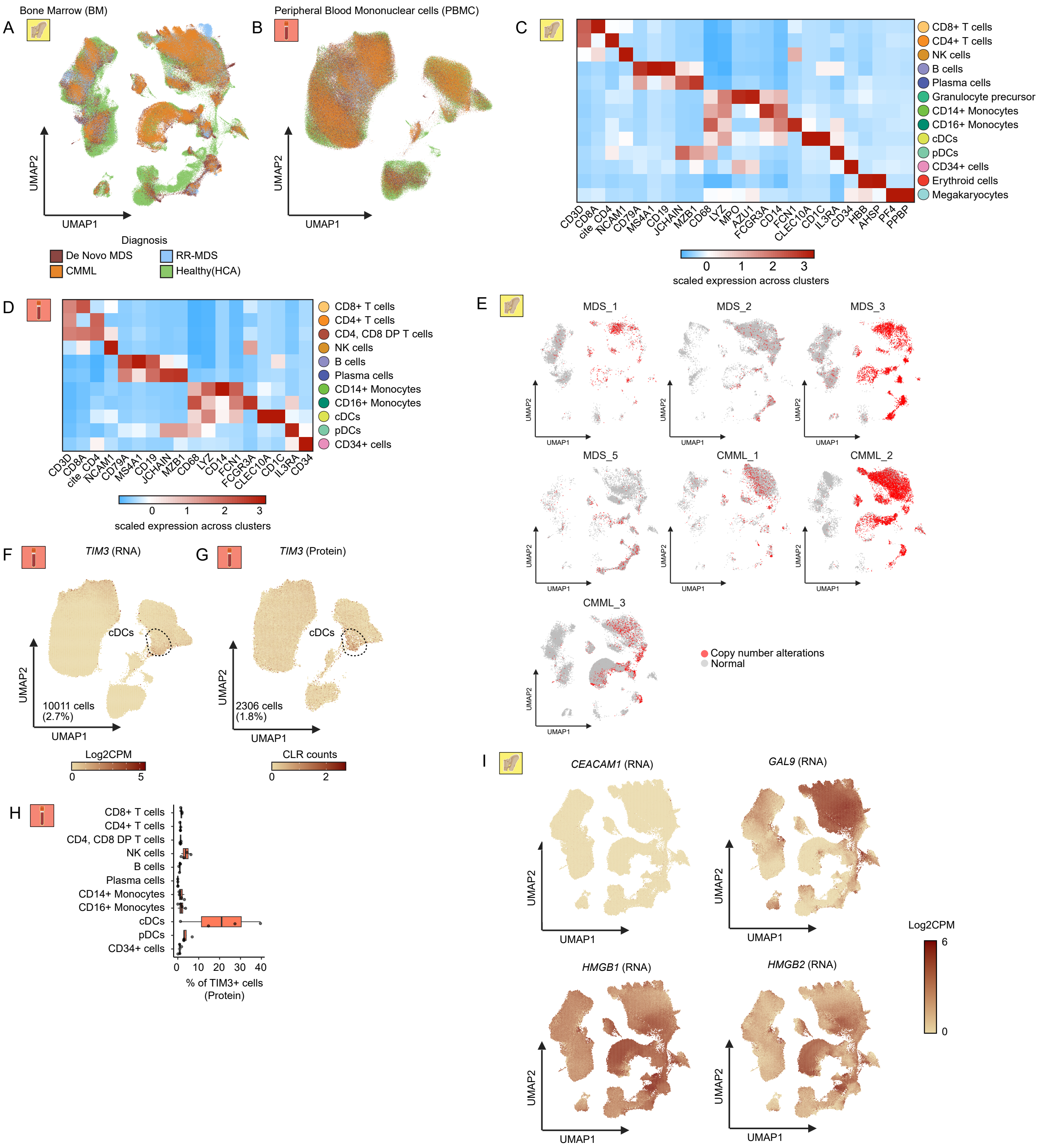

A
Bone Marrow (BM)
UMAP1
UMAP2
B
Peripheral Blood Mononuclear cells (PBMC)
Diagnosis
De Novo MDS
RR-MDS
CMML
Healthy(HCA)
C
CD8+ T cells
CD4+ T cells
NK cells
B cells
Plasma cells
Granulocyte precursor
CD14+ Monocytes
CD16+ Monocytes
cDCs
pDCs
CD34+ cells
Erythroid cells
Megakaryocytes
CD3D CD8A cite_CD4 NCAM1 CD79A MS4A1 CD19 JCHAIN MZB1 CD68 LYZ MPO AZU1 FCGR3A CD14 FCN1 CLEC10A CD1C IL3RA CD34 HBB AHSP PF4 PPBP
scaled expression across clusters
D
CD4, CD8 DP T cells
E
MDS_1
MDS_2
MDS_3
MDS_5
CMML_1
CMML_2
CMML_3
Copy number alterations
Normal
F
TIM3 (RNA)
cDCs
10011 cells (2.7%)
Log2CPM
G
TIM3 (Protein)
2306 cells (1.8%)
CLR counts
H
% of TIM3+ cells (Protein)
I
CEACAM1 (RNA)
GAL9 (RNA)
HMGB1 (RNA)
HMGB2 (RNA)

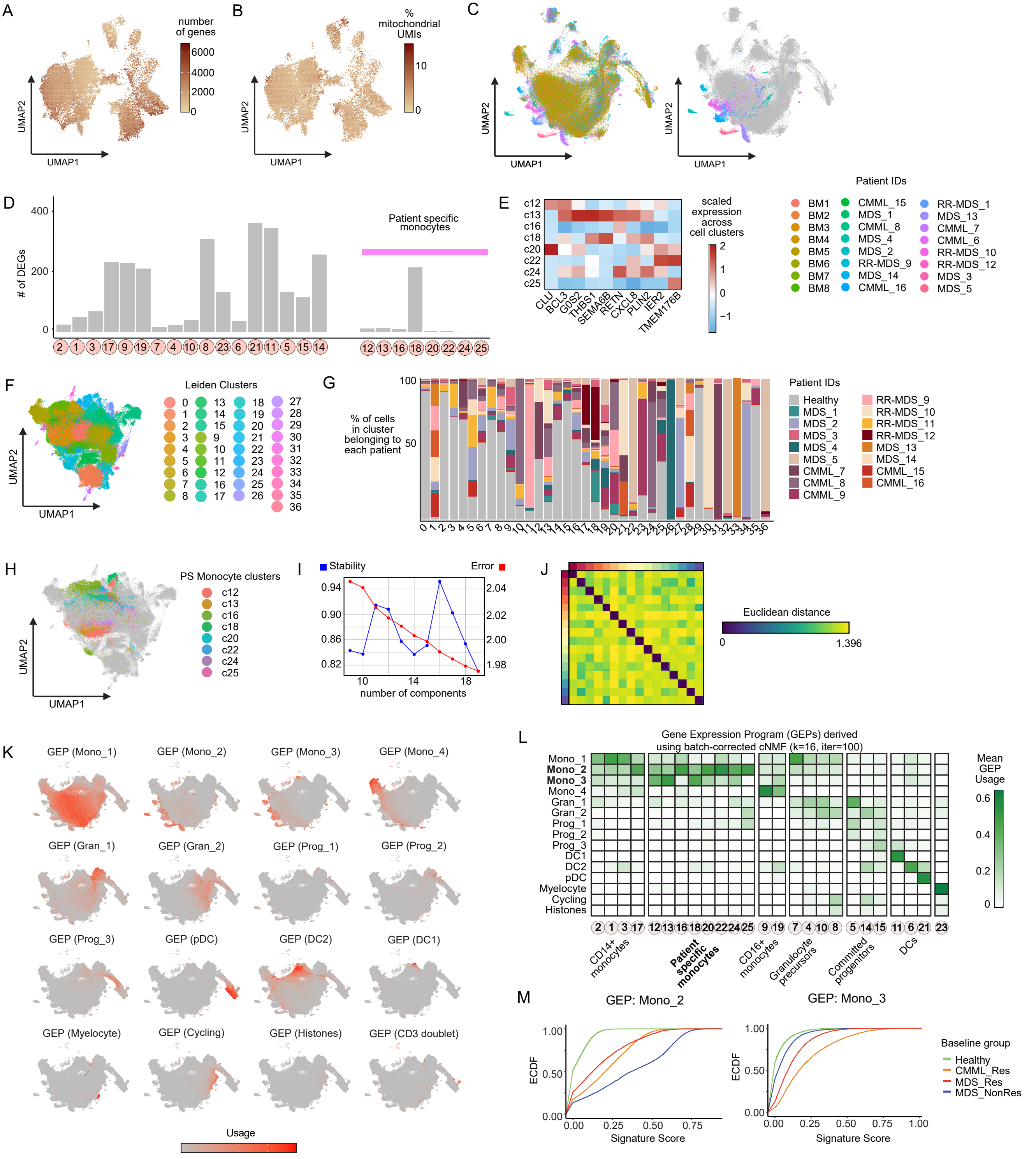
A
number of genes
6000
4000
2000
0
UMAP2
UMAP1
B
% mitochondrial UMIs
10
0
UMAP2
UMAP1
C
UMAP2
UMAP1
UMAP2
UMAP1
Patient IDs
BM1
BM2
BM3
BM4
BM5
BM6
BM7
BM8
CMML_15
MDS_1
CMML_8
MDS_4
MDS_2
RR-MDS_9
MDS_14
CMML_16
RR-MDS_1
MDS_13
CMML_7
CMML_6
RR-MDS_10
RR-MDS_12
MDS_3
MDS_5
D
### of DEGs
400
200
0
Patient specific monocytes
2 1 3 17 9 19 7 4 10 8 23 6 21 11 5 15 14
12 13 16 18 20 22 24 25
E
c12
c13
c16
c18
c20
c22
c24
c25
CLU
BCL3
G0S2
THBS1
SEMA6B
RETN
CXCL8
PLIN2
IER2
TMEM176B
scaled expression across cell clusters
2
1
0
−1
F
Leiden Clusters
UMAP2
UMAP1
G
% of cells in cluster belonging to each patient
100
50
Patient IDs
Healthy
MDS_1
MDS_2
MDS_3
MDS_4
MDS_5
CMML_7
CMML_8
CMML_9
RR-MDS_9
RR-MDS_10
RR-MDS_11
RR-MDS_12
MDS_13
MDS_14
CMML_15
CMML_16
H
PS Monocyte clusters
c12
c13
c16
c18
c20
c22
c24
c25
UMAP2
UMAP1
I
Stability
Error
0.94
0.90
0.86
0.82
2.04
2.02
2.00
1.98
10
14
18
number of components
J
Euclidean distance
0
1.396
K
GEP (Mono_1)
GEP (Mono_2)
GEP (Mono_3)
GEP (Mono_4)
GEP (Gran_1)
GEP (Gran_2)
GEP (Prog_1)
GEP (Prog_2)
GEP (Prog_3)
GEP (pDC)
GEP (DC2)
GEP (DC1)
GEP (Myelocyte)
GEP (Cycling)
GEP (Histones)
GEP (CD3 doublet)
Usage
L
Gene Expression Program (GEPs) derived using batch-corrected cNMF (k=16, iter=100)
Mono_1
Mono_2
Mono_3
Mono_4
Gran_1
Gran_2
Prog_1
Prog_2
Prog_3
DC1
DC2
pDC
Myelocyte
Cycling
Histones
Mean GEP Usage
0.6
0.4
0.2
0
2 1 3 17
12 13 16 18 20 22 24 25
9 19
7 4 10 8
5 14 15
11 6 21
23
CD14+ monocytes
Patient specific monocytes
CD16+ monocytes
Granulocyte precursors
Committed progenitors
DCs
M
GEP: Mono_2
ECDF
1.00
0.50
0.00
0.00
0.25
0.50
0.75
Signature Score
GEP: Mono_3
ECDF
1.00
0.50
0.00
0.00
0.25
0.50
0.75
1.00
Signature Score
Baseline group
Healthy
CMML_Res
MDS_Res
MDS_NonRes

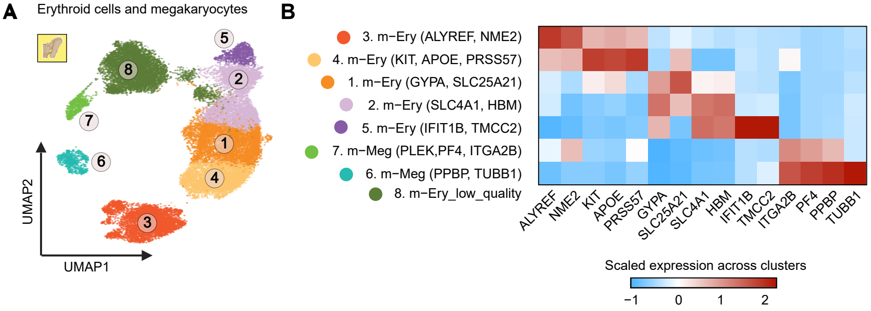
A
Erythroid cells and megakaryocytes
UMAP2
UMAP1
B
3. m−Ery (ALYREF, NME2)
4. m−Ery (KIT, APOE, PRSS57)
1. m−Ery (GYPA, SLC25A21)
2. m−Ery (SLC4A1, HBM)
5. m−Ery (IFIT1B, TMCC2)
7. m−Meg (PLEK,PF4, ITGA2B)
6. m−Meg (PPBP, TUBB1)
8. m−Ery_low_quality
ALYREF
NME2
KIT
APOE
PRSS57
GYPA
SLC25A21
SLC4A1
HBM
IFIT1B
TMCC2
ITGA2B
PF4
PPBP
TUBB1
Scaled expression across clusters
−1
0
1
2

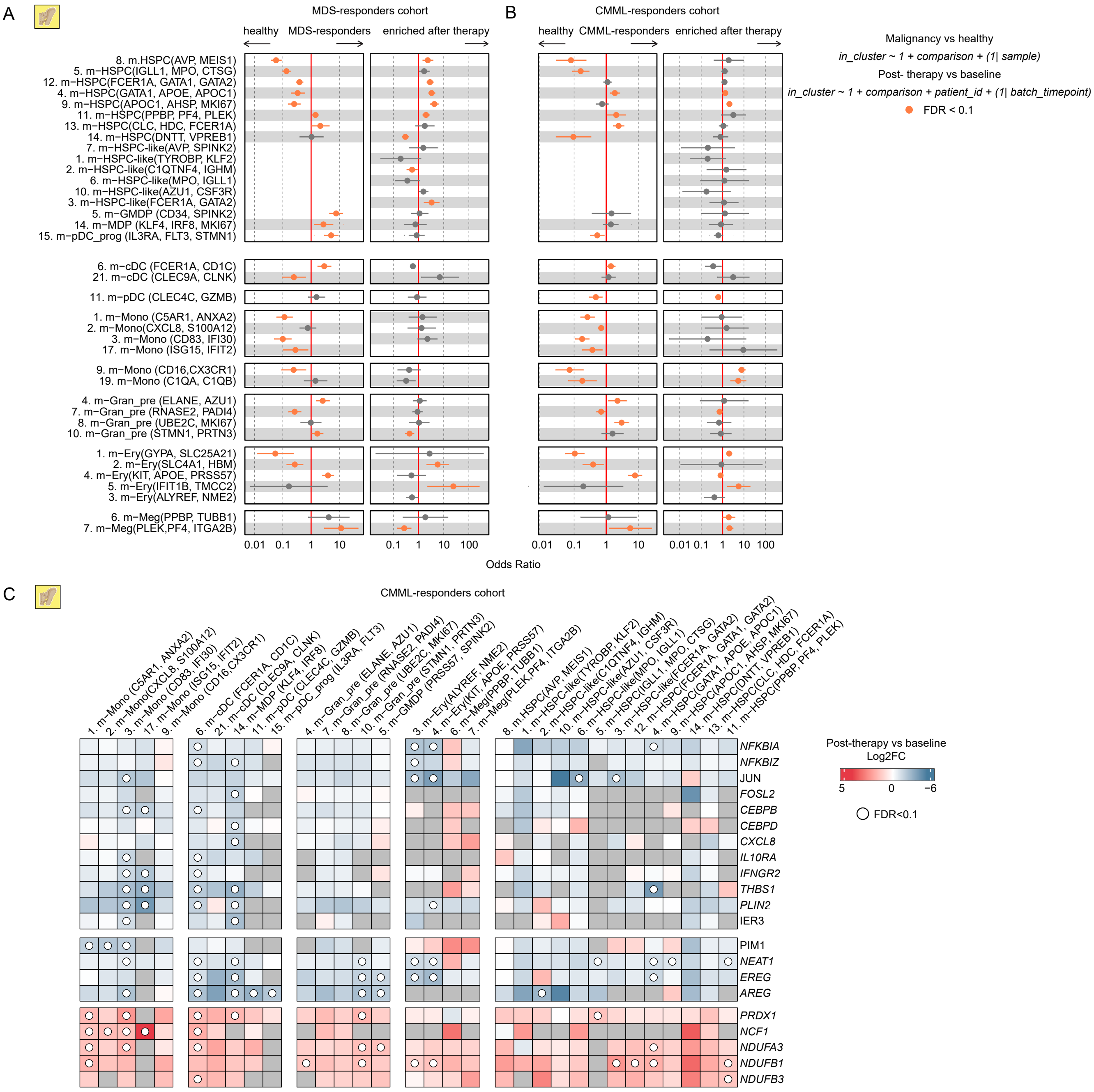

A
MDS-responders cohort
healthy
MDS-responders
enriched after therapy
B
CMML-responders cohort
healthy
CMML-responders
enriched after therapy
Malignancy vs healthy
in_cluster ~ 1 + comparison + (1| sample)
Post- therapy vs baseline
in_cluster ~ 1 + comparison + patient_id + (1| batch_timepoint)
FDR < 0.1
8. m.HSPC(AVP, MEIS1)
5. m−HSPC(IGLL1, MPO, CTSG)
12. m−HSPC(FCER1A, GATA1, GATA2)
4. m−HSPC(GATA1, APOE, APOC1)
9. m−HSPC(APOC1, AHSP, MKI67)
11. m−HSPC(PPBP, PF4, PLEK)
13. m−HSPC(CLC, HDC, FCER1A)
14. m−HSPC(DNTT, VPREB1)
7. m−HSPC-like(AVP, SPINK2)
1. m−HSPC-like(TYROBP, KLF2)
2. m−HSPC-like(C1QTNF4, IGHM)
6. m−HSPC-like(MPO, IGLL1)
10. m−HSPC-like(AZU1, CSF3R)
3. m−HSPC-like(FCER1A, GATA2)
5. m−GMDP (CD34, SPINK2)
14. m−MDP (KLF4, IRF8, MKI67)
15. m−pDC_prog (IL3RA, FLT3, STMN1)
6. m−cDC (FCER1A, CD1C)
21. m−cDC (CLEC9A, CLNK)
11. m−pDC (CLEC4C, GZMB)
1. m−Mono (C5AR1, ANXA2)
2. m−Mono(CXCL8, S100A12)
3. m−Mono (CD83, IFI30)
17. m−Mono (ISG15, IFIT2)
9. m−Mono (CD16,CX3CR1)
19. m−Mono (C1QA, C1QB)
4. m−Gran_pre (ELANE, AZU1)
7. m−Gran_pre (RNASE2, PADI4)
8. m−Gran_pre (UBE2C, MKI67)
10. m−Gran_pre (STMN1, PRTN3)
1. m−Ery(GYPA, SLC25A21)
2. m−Ery(SLC4A1, HBM)
4. m−Ery(KIT, APOE, PRSS57)
5. m−Ery(IFIT1B, TMCC2)
3. m−Ery(ALYREF, NME2)
6. m−Meg(PPBP, TUBB1)
7. m−Meg(PLEK,PF4, ITGA2B)
0.01 0.1 1 10
0.1 1 10 100
0.01 0.1 1 10
0.01 0.1 1 10 100
Odds Ratio
C
CMML-responders cohort
1. m-Mono (C5AR1, ANXA2)
2. m-Mono(CXCL8, S100A12)
3. m-Mono (CD83, IFI30)
17. m-Mono (ISG15, IFIT2)
9. m-Mono (CD16,CX3CR1)
6. m-cDC (FCER1A, CD1C)
21. m-cDC (CLEC9A, CLNK)
14. m-MDP (KLF4, IRF8)
11. m-pDC (CLEC4C, GZMB)
15. m-pDC_prog (IL3RA, FLT3)
4. m-Gran_pre (ELANE, AZU1)
7. m-Gran_pre (RNASE2, PADI4)
8. m-Gran_pre (UBE2C, MKI67)
10. m-Gran_pre (STMN1, PRTN3)
5. m-GMDP (PRSS57, SPINK2)
3. m-Ery(ALYREF, NME2)
4. m-Ery(KIT, APOE, PRSS57)
6. m-Meg(PPBP, TUBB1)
7. m-Meg(PLEK,PF4, ITGA2B)
8. m.HSPC(AVP, MEIS1)
1. m-HSPC-like(TYROBP, KLF2)
2. m-HSPC-like(C1QTNF4, IGHM)
10. m-HSPC-like(AZU1, CSF3R)
6. m-HSPC-like(MPO, IGLL1)
5. m-HSPC(IGLL1, MPO, CTSG)
3. m-HSPC-like(FCER1A, GATA2)
12. m-HSPC(FCER1A, GATA1, GATA2)
4. m-HSPC(GATA1, APOE, APOC1)
9. m-HSPC(APOC1, AHSP, MKI67)
14. m-HSPC(DNTT, VPREB1)
13. m-HSPC(CLC, HDC, FCER1A)
11. m-HSPC(PPBP, PF4, PLEK)
NFKBIA
NFKBIZ
JUN
FOSL2
CEBPB
CEBPD
CXCL8
IL10RA
IFNGR2
THBS1
PLIN2
IER3
PIM1
NEAT1
EREG
AREG
PRDX1
NCF1
NDUFA3
NDUFB1
NDUFB3
Post-therapy vs baseline
Log2FC
5
0
−6
FDR<0.1

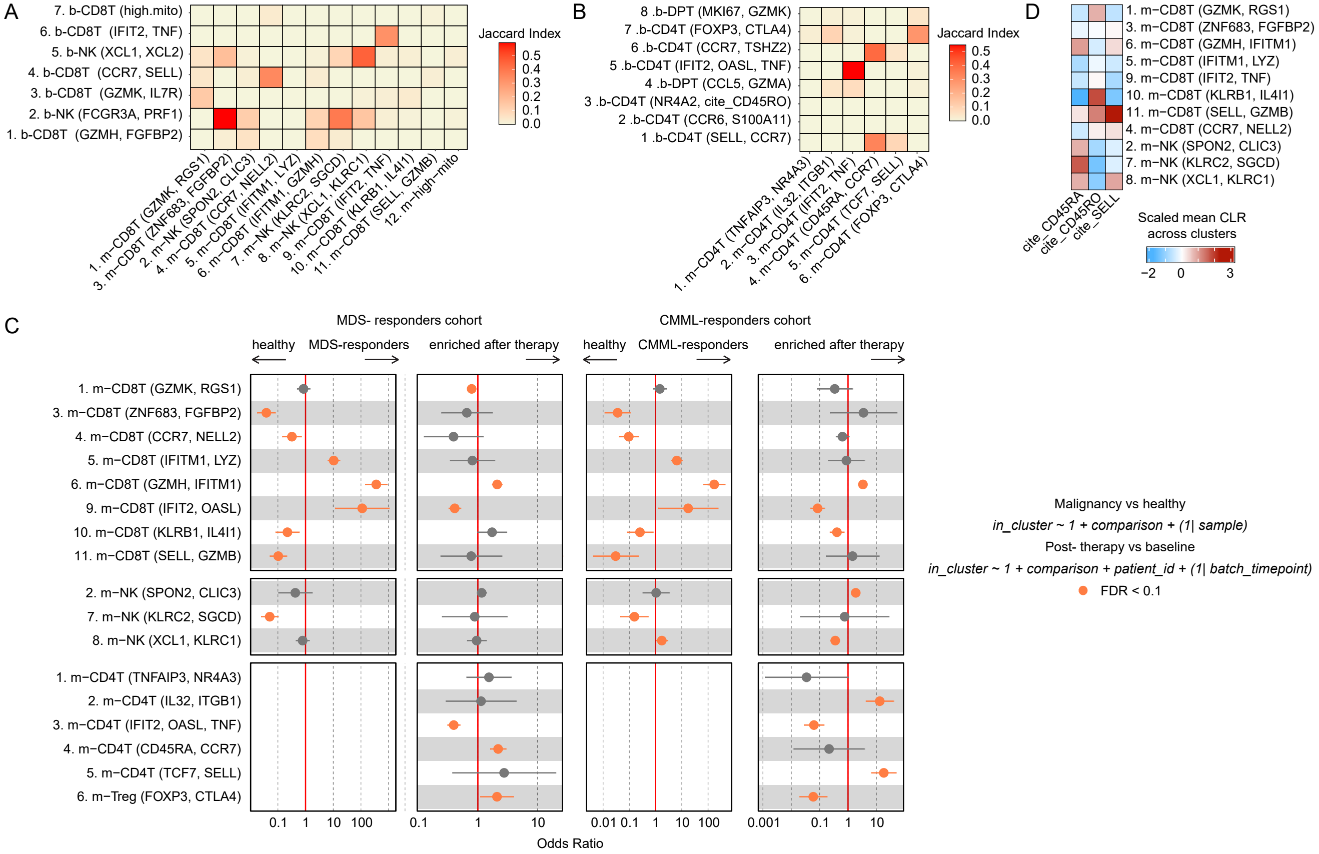

A
7. b-CD8T (high.mito)
6. b-CD8T (IFIT2, TNF)
5. b-NK (XCL1, XCL2)
4. b-CD8T (CCR7, SELL)
3. b-CD8T (GZMK, IL7R)
2. b-NK (FCGR3A, PRF1)
1. b-CD8T (GZMH, FGFBP2)
1. m−CD8T (GZMK, RGS1)
3. m−CD8T (ZNF683, FGFBP2)
2. m−NK (SPON2, CLIC3)
4. m−CD8T (CCR7, NELL2)
5. m−CD8T (IFITM1, LYZ)
6. m−CD8T (IFITM1, GZMH)
7. m−NK (KLRC2, SGCD)
8. m−NK (XCL1, KLRC1)
9. m−CD8T (IFIT2, TNF)
10. m−CD8T (KLRB1, IL4I1)
11. m−CD8T (SELL, GZMB)
12. m−high−mito
Jaccard Index
0.5
0.4
0.3
0.2
0.1
0.0
B
8 .b-DPT (MKI67, GZMK)
7 .b-CD4T (FOXP3, CTLA4)
6 .b-CD4T (CCR7, TSHZ2)
5 .b-CD4T (IFIT2, OASL, TNF)
4 .b-DPT (CCL5, GZMA)
3 .b-CD4T (NR4A2, cite_CD45RO)
2 .b-CD4T (CCR6, S100A11)
1 .b-CD4T (SELL, CCR7)
1. m−CD4T (TNFAIP3, NR4A3)
2. m−CD4T (IL32, ITGB1)
3. m−CD4T (IFIT2, TNF)
4. m−CD4T (CD45RA, CCR7)
5. m−CD4T (TCF7, SELL)
6. m−CD4T (FOXP3, CTLA4)
Jaccard Index
0.5
0.4
0.3
0.2
0.1
0.0
D
1. m−CD8T (GZMK, RGS1)
3. m−CD8T (ZNF683, FGFBP2)
6. m−CD8T (GZMH, IFITM1)
5. m−CD8T (IFITM1, LYZ)
9. m−CD8T (IFIT2, TNF)
10. m−CD8T (KLRB1, IL4I1)
11. m−CD8T (SELL, GZMB)
4. m−CD8T (CCR7, NELL2)
2. m−NK (SPON2, CLIC3)
7. m−NK (KLRC2, SGCD)
8. m−NK (XCL1, KLRC1)
cite_CD45RA
cite_CD45RO
cite_SELL
Scaled mean CLR across clusters
−2
0
3
C
MDS- responders cohort
CMML-responders cohort
healthy
MDS-responders
enriched after therapy
healthy
CMML-responders
enriched after therapy
1. m−CD8T (GZMK, RGS1)
3. m−CD8T (ZNF683, FGFBP2)
4. m−CD8T (CCR7, NELL2)
5. m−CD8T (IFITM1, LYZ)
6. m−CD8T (GZMH, IFITM1)
9. m−CD8T (IFIT2, OASL)
10. m−CD8T (KLRB1, IL4I1)
11. m−CD8T (SELL, GZMB)
2. m−NK (SPON2, CLIC3)
7. m−NK (KLRC2, SGCD)
8. m−NK (XCL1, KLRC1)
1. m−CD4T (TNFAIP3, NR4A3)
2. m−CD4T (IL32, ITGB1)
3. m−CD4T (IFIT2, OASL, TNF)
4. m−CD4T (CD45RA, CCR7)
5. m−CD4T (TCF7, SELL)
6. m−Treg (FOXP3, CTLA4)
0.1
1
10
100
0.1
1
10
0.01
0.1
1
10
100
0.001
0.1
1
10
Odds Ratio
Malignancy vs healthy
in_cluster ~ 1 + comparison + (1| sample)
Post- therapy vs baseline
in_cluster ~ 1 + comparison + patient_id + (1| batch_timepoint)
FDR < 0.1

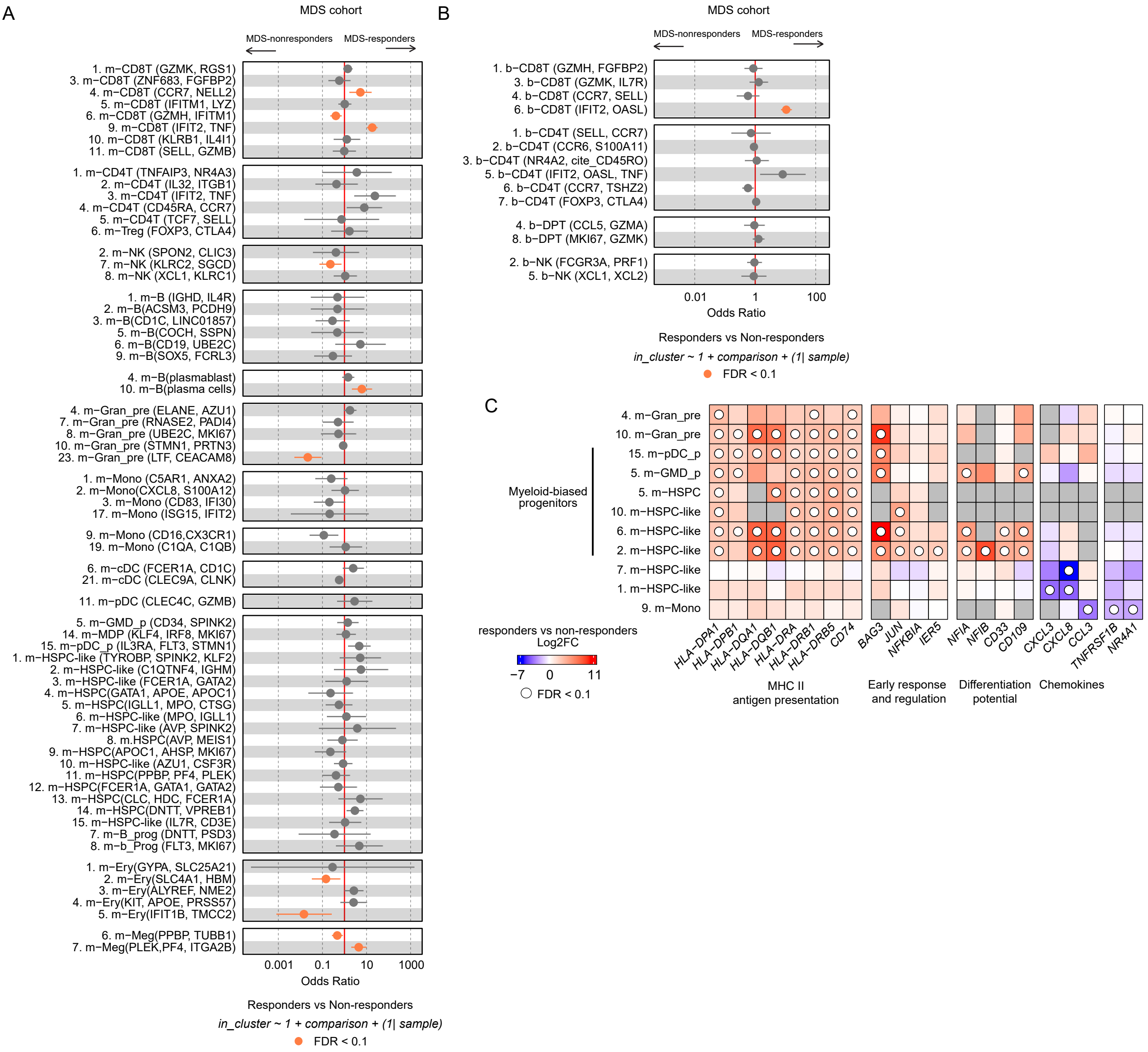
A
MDS cohort
MDS-nonresponders
MDS-responders
1. m−CD8T (GZMK, RGS1)
3. m−CD8T (ZNF683, FGFBP2)
4. m−CD8T (CCR7, NELL2)
5. m−CD8T (IFITM1, LYZ)
6. m−CD8T (GZMH, IFITM1)
9. m−CD8T (IFIT2, TNF)
10. m−CD8T (KLRB1, IL4I1)
11. m−CD8T (SELL, GZMB)
1. m−CD4T (TNFAIP3, NR4A3)
2. m−CD4T (IL32, ITGB1)
3. m−CD4T (IFIT2, TNF)
4. m−CD4T (CD45RA, CCR7)
5. m−CD4T (TCF7, SELL)
6. m−Treg (FOXP3, CTLA4)
2. m−NK (SPON2, CLIC3)
7. m−NK (KLRC2, SGCD)
8. m−NK (XCL1, KLRC1)
1. m−B (IGHD, IL4R)
2. m−B(ACSM3, PCDH9)
3. m−B(CD1C, LINC01857)
5. m−B(COCH, SSPN)
6. m−B(CD19, UBE2C)
9. m−B(SOX5, FCRL3)
4. m−B(plasmablast)
10. m−B(plasma cells)
4. m−Gran_pre (ELANE, AZU1)
7. m−Gran_pre (RNASE2, PADI4)
8. m−Gran_pre (UBE2C, MKI67)
10. m−Gran_pre (STMN1, PRTN3)
23. m−Gran_pre (LTF, CEACAM8)
1. m−Mono (C5AR1, ANXA2)
2. m−Mono(CXCL8, S100A12)
3. m−Mono (CD83, IFI30)
17. m−Mono (ISG15, IFIT2)
9. m−Mono (CD16,CX3CR1)
19. m−Mono (C1QA, C1QB)
6. m−cDC (FCER1A, CD1C)
21. m−cDC (CLEC9A, CLNK)
11. m−pDC (CLEC4C, GZMB)
5. m−GMD_p (CD34, SPINK2)
14. m−MDP (KLF4, IRF8, MKI67)
15. m−pDC_p (IL3RA, FLT3, STMN1)
1. m−HSPC-like (TYROBP, SPINK2, KLF2)
2. m−HSPC-like (C1QTNF4, IGHM)
3. m−HSPC-like (FCER1A, GATA2)
4. m−HSPC(GATA1, APOE, APOC1)
5. m−HSPC(IGLL1, MPO, CTSG)
6. m−HSPC-like (MPO, IGLL1)
7. m−HSPC-like (AVP, SPINK2)
8. m.HSPC(AVP, MEIS1)
9. m−HSPC(APOC1, AHSP, MKI67)
10. m−HSPC-like (AZU1, CSF3R)
11. m−HSPC(PPBP, PF4, PLEK)
12. m−HSPC(FCER1A, GATA1, GATA2)
13. m−HSPC(CLC, HDC, FCER1A)
14. m−HSPC(DNTT, VPREB1)
15. m−HSPC-like (IL7R, CD3E)
7. m−B_prog (DNTT, PSD3)
8. m−b_Prog (FLT3, MKI67)
1. m−Ery(GYPA, SLC25A21)
2. m−Ery(SLC4A1, HBM)
3. m−Ery(ALYREF, NME2)
4. m−Ery(KIT, APOE, PRSS57)
5. m−Ery(IFIT1B, TMCC2)
6. m−Meg(PPBP, TUBB1)
7. m−Meg(PLEK,PF4, ITGA2B)
0.001
0.1
10
1000
Odds Ratio
Responders vs Non-responders
in_cluster ~ 1 + comparison + (1| sample)
FDR < 0.1
B
MDS cohort
MDS-nonresponders
MDS-responders
1. b−CD8T (GZMH, FGFBP2)
3. b−CD8T (GZMK, IL7R)
4. b−CD8T (CCR7, SELL)
6. b−CD8T (IFIT2, OASL)
1. b−CD4T (SELL, CCR7)
2. b−CD4T (CCR6, S100A11)
3. b−CD4T (NR4A2, cite_CD45RO)
5. b−CD4T (IFIT2, OASL, TNF)
6. b−CD4T (CCR7, TSHZ2)
7. b−CD4T (FOXP3, CTLA4)
4. b−DPT (CCL5, GZMA)
8. b−DPT (MKI67, GZMK)
2. b−NK (FCGR3A, PRF1)
5. b−NK (XCL1, XCL2)
0.01
1
100
Odds Ratio
Responders vs Non-responders
in_cluster ~ 1 + comparison + (1| sample)
FDR < 0.1
C
4. m−Gran_pre
10. m−Gran_pre
15. m−pDC_p
5. m−GMD_p
5. m−HSPC
10. m−HSPC-like
6. m−HSPC-like
2. m−HSPC-like
7. m−HSPC-like
1. m−HSPC-like
9. m−Mono
Myeloid-biased progenitors
HLA-DPA1
HLA-DPB1
HLA-DQA1
HLA-DQB1
HLA-DRA
HLA-DRB1
HLA-DRB5
CD74
BAG3
JUN
NFKBIA
IER5
NFIA
NFIB
CD33
CD109
CXCL3
CXCL8
CCL3
TNFRSF1B
NR4A1
MHC II antigen presentation
Early response and regulation
Differentiation potential
Chemokines
responders vs non-responders
Log2FC
−7
0
11
FDR < 0.1

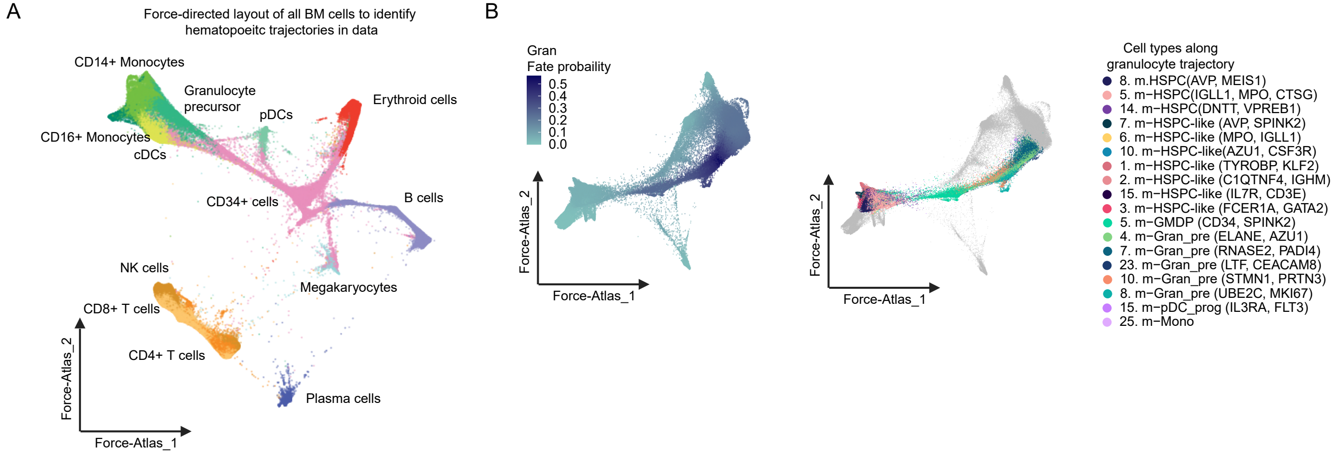

A
Force-directed layout of all BM cells to identify hematopoeitc trajectories in data
CD14+ Monocytes
Granulocyte precursor
pDCs
Erythroid cells
CD16+ Monocytes
cDCs
CD34+ cells
B cells
NK cells
Megakaryocytes
CD8+ T cells
CD4+ T cells
Plasma cells
Force-Atlas_2
Force-Atlas_1
B
Gran
Fate probaility
0.5
0.4
0.3
0.2
0.1
0.0
Force-Atlas_2
Force-Atlas_1
Force-Atlas_2
Force-Atlas_1
Cell types along granulocyte trajectory
8. m.HSPC(AVP, MEIS1)
5. m−HSPC(IGLL1, MPO, CTSG)
14. m−HSPC(DNTT, VPREB1)
7. m−HSPC-like (AVP, SPINK2)
6. m−HSPC-like (MPO, IGLL1)
10. m−HSPC-like(AZU1, CSF3R)
1. m−HSPC-like (TYROBP, KLF2)
2. m−HSPC-like (C1QTNF4, IGHM)
15. m−HSPC-like (IL7R, CD3E)
3. m−HSPC-like (FCER1A, GATA2)
5. m−GMDP (CD34, SPINK2)
4. m−Gran_pre (ELANE, AZU1)
7. m−Gran_pre (RNASE2, PADI4)
23. m−Gran_pre (LTF, CEACAM8)
10. m−Gran_pre (STMN1, PRTN3)
8. m−Gran_pre (UBE2C, MKI67)
15. m−pDC_prog (IL3RA, FLT3)
25. m−Mono
