## Supplementary Figure Legends for "Response to azacitidine and the TIM-3 inhibitor, sabatolimab, in MDS and CMML is associated with distinct immune and antigen-presentation states"

**Supplementary Figures**

**Supplementary Figure 1. Major lineage annotation markers, inferred copy number alterations, and TIM3 and ligand expression across bone marrow and peripheral blood. A-B.** UMAP embedding of (**A**) 379,284 cells from bone marrow (BM) and (**B**) 367,973 cells from peripheral blood (PB) colored by sample origin group. **C.** Heatmap showing select discriminative markers for each major lineage in **A** when compared with every other lineage. **D.** Heatmap showing select discriminative markers for each major lineage in **B** when compared with every other lineage. **E.** UMAP embeddings of BM cells from individual patients colored by NUMBAT inferred copy number alteration (CNA) status. **F-G**. UMAP feature plots using color to indicate RNA transcript (**F**; log2CPM) and protein (**G**; CLR) levels of *HAVCR2*/TIM3 in PBMC. Number and percentage correspond to cells with non-zero expression. **H.** Percent of cells from individual samples in each lineage with non-zero counts for TIM3 protein after background signature removal. Box plots show median (line) and IQR, with whiskers no more than 1.5x IQR. **I.** UMAP feature plots using color to indicate RNA transcript (log2CPM) levels of *CEACAM1*, *GAL9*, *HMGB1*, and *HMGB2*. In **C** and **D,** Color scale denotes the normalized gene expression (mean zero, unit variance) across lineages.

**Supplementary Figure 2. Quality-control metrics, batch correction comparison, and cNMF gene expression program derivation for myelomonocytic and dendritic cells.**

**A.** UMAP feature plots showing number of genes  **B.** UMAP feature plots showing percentage of mitochondrial transcripts in CD34 cell embedding. **C.** UMAP embedding of myelomonocytic cells and dendritic cells (DCs) from BM colored by patient ID (left) and patient ID within patient-specific monocyte subsets only (right). **D.** Barplot showing the number of differentially expressed genes (DEGs) when comparing each cluster to all other clusters. DEGs are defined as genes with Log2FoldChange > 1 and adjusted P value < 0.05 (limma). **E.** Heatmap showing select discriminative markers for each patient specific monocyte cluster when compared with every other cluster. Color scale denotes the normalized gene expression (mean zero, unit variance) across cell clusters. **F.** UMAP embedding of myelomonocytic cells and DCs from BM samples, generated using the latent representation learned using scVI for dimensionality reduction, colored by new clusters. **G.** Stacked barplot showing cell origin composition of new clusters, with all healthy donors collapsed into a single category. **H.** UMAP embedding of myelomonocytic and DCs from BM samples learned using scVI, colored by patient specific monocyte subsets identified from the Harmony-corrected representation.  **I.** Line chart showing the stability and reconstruction error across a range of gene expression programs(GEPs) (k). **J.** Heatmap showing the Euclinean distance between all factors grouped together to generate k=16 GEPs. **K.** Feature plots of the GEP usage in myelomonocytic cells and DCs from BM. **L.** Heatmap showing mean usage of GEPs across all myelomonocytic and DC clusters. **M.** Empirical Cumulative Distribution Function (ECDF) of Mono_2 (left) and Mono_3 (right) GEP signature scores in myelomonocytic and DC populations split by baseline sample grouping. At a given signature score, ECDF=1 indicates all cells have a signature score below said score and an ECDF=0.5 indicates half the cells have a score higher than said score.

**Supplementary Figure 3. Overview of erythroid cells and megakaryocytes annotation markers.**

**A.**  UMAP embedding of 26,523 erythroid cells and megakaryocytes  from BM colored by subcluster annotations. **B.** Heatmap showing select discriminative markers for each cell cluster when compared with every other cluster.

**Supplementary Figure 4. Post-therapy abundance shifts and DEGs in myelomonocytic and dendritic cells,** **and progenitor cells in MDS and CMML responders.**

**A.** Cell cluster abundance analysis in the BM comparing MDS-responders to healthy donors at baseline (left), and comparing MDS-responders post-treatment to pre-treatment samples (right). **B.** Cell cluster abundance analysis in the BM comparing CMML-responders to healthy donors at baseline (left), and comparing CMML-responders post-treatment to pre-treatment samples (right). **C.** Heatmap showing select DEGs in the BM among CMML-responders for all myeloid, erythroid and progenitor cell subsets comparing post-treatment to pre-treatment samples. Color scale indicates log2FC and white dots indicate FDR < 0.1. In **A** and **B,** Points represent odds-ratio with error bars representing 95% confidence intervals. Color indicates FDR < 0.1 (logistic regression).

**Supplementary Figure 5. Overview of CD4 T, CD8 T and NK cell annotation markers and post-therapy abundance shifts in these cell populations in MDS and CMML responders.**

**A.** Heatmap showing overlap of transcriptionally defined marker genes between CD8 T cell subsets in BM and PBMC. Color indicates Jaccard Index for each pair of cell subsets across the two tissue compartments. **B.** Heatmap showing overlap of transcriptionally defined marker genes between CD4 T cell subsets in BM and PBMC. Color indicates Jaccard Index for each pair of cell subsets across the two tissue compartments. **C.** Heatmap showing protein expression of CD45RA, CD45RO and SELL for each CD8 T and NK cell cluster when compared with every other cluster.  **D.** Cell cluster abundance analysis comparing MDS-responders to healthy donors at baseline (left most panel); comparing MDS-responders post-treatment to pre-treatment samples (second to left panel); comparing CMML-responders to healthy donors at baseline (second to right panel); and comparing CMML-responders post-treatment to pre-treatment samples (right most panel).

**Supplementary Figure 6. Baseline abundance shifts and DEGs between MDS-responder and non-responders.**

**A.** Cell cluster abundance analysis for all annotated cell subsets in the BM comparing the baseline timepoint from MDS-responders and MDS-non-responders. **B.** Cell cluster abundance analysis for CD8 T cell, NK cell and CD4 T cell subsets in PBMC comparing MDS-responders to MDS-non-responders at baseline. **C.** Heatmap showing select DEGs for select hematopoietic stem cell, progenitor and myeloid precursor cell subsets in BM when comparing MDS-responders to MDS-non-responders at baseline. Color scale indicates log2FC and white dots indicate FDR < 0.1. In **A** and **B,** Points represent odds-ratio with error bars representing 95% confidence intervals. Color indicates FDR < 0.1 (logistic regression).

**Supplementary Figure 7. Identification of granulocyte trajectory from lineage map of all bone marrow cells.**

**A.** Force-Atlas2 embedding of all cells from BM colored by major cell lineage. **B.** Force-Atlas2 embedding of myelomonocytic and dendritic cells, and HSPCs from BM colored by inferred granulocyte fate probability (left), and original cell subset annotations of cells within granulocyte trajectory (right).
